# An IRF4-MYC-mTORC1 integrated pathway controls cell growth and the proliferative capacity of activated B cells during B cell differentiation *in vivo*

**DOI:** 10.1101/2021.06.20.449155

**Authors:** Dillon G. Patterson, Anna K. Kania, Madeline J. Price, James R. Rose, Christopher D. Scharer, Jeremy M. Boss

## Abstract

Cell division is an essential component of B cell differentiation to antibody-secreting plasma cells, with critical reprogramming occurring during the initial stages of B cell activation. However, a complete understanding of the factors that coordinate early reprogramming events in vivo remain to be determined. In this study, we examined the initial reprogramming by IRF4 in activated B cells using an adoptive transfer system and mice with a B cell-specific deletion of IRF4. IRF4-deficient B cells responding to influenza, NP-Ficoll and LPS divided, but stalled during the proliferative response. Gene expression profiling of IRF4-deficient B cells at discrete divisions revealed IRF4 was critical for inducing MYC target genes, oxidative phosphorylation, and glycolysis. Moreover, IRF4-deficient B cells maintained an inflammatory gene expression signature. Complementary chromatin accessibility analyses established a hierarchy of IRF4 activity and identified networks of dysregulated transcription factor families in IRF4-deficient B cells, including E-box binding bHLH family members. Indeed, B cells lacking IRF4 failed to fully induce *Myc* after stimulation and displayed aberrant cell cycle distribution. Furthermore, IRF4-deficient B cells showed reduced mTORC1 activity and failed to initiate the B cell-activation unfolded protein response and grow in cell size. *Myc* overexpression in IRF4-deficient was sufficient to overcome the cell growth defect. Together, these data reveal an IRF4-MYC-mTORC1 relationship critical for controlling cell growth and the proliferative response during B cell differentiation.

## Introduction

A key component of the adaptive immune response is the generation of antibody by antibody-secreting plasma cells (ASC). Upon antigen encounter, quiescent naïve B cells become activated, rapidly proliferate, and a subset differentiate to ASC. One essential component of B cell differentiation to ASC is cell division (1-4). Culturing purified B cells and blocking cell division prevents the generation of ASC (3). However, the number of cell divisions does not exclusively determine ASC formation. This has led to a stochastic model of differentiation that describes population-level immune responses and accounts for heterogeneity in cell fates among responding cells, such as whether they will continue to divide, die, or differentiate (1, 5-7). One molecular determinant that contributes to such heterogeneity is the expression levels of MYC (4, 8-12). MYC levels are influenced by immune stimulation and serve as a division-independent timer to control the proliferative capacity of responding cells (4, 10, 13). IRF4 is another factor that contributes to heterogeneity at the population level (14-17). During the initial stages of B cell activation, high IRF4 expression biases cells towards the ASC fate (14, 17). Notably, initial IRF4 expression levels are influenced by the intensity of immune stimulation, and IRF4^hi^ cells are among the first to divide (18). Indeed, proliferation is reduced in IRF4-deficient B cells stimulated ex vivo (15, 16, 19); however, the impact of IRF4 on in vivo B cell proliferation is unknown. Furthermore, the timing, scope, and mechanism by which IRF4 contributes to control the proliferative response remains undefined.

Cell division is tightly linked to ASC formation, with transcriptional and epigenetic reprogramming (20-23) occurring as the cells divide (17, 24-26). As such, each cellular division represents a distinct stage during B cell differentiation, with ASC formation occurring after at least eight cell divisions (17, 25, 27). Cell extrinsic signals can impact the specific division in which differentiation occurs, but the molecular programming events leading to ASC remain the same (17). Many essential ASC programming events (28) are initiated during the early stages of B cell activation and are progressively reinforced in subsequent divisions (24, 25, 27). For example, ASC formation requires a metabolic shift from glycolysis to oxidative phosphorylation (OXPHOS), and the OXPHOS program is increasingly established across cell divisions (24). Additionally, ASC differentiation requires activation of the unfolded protein response (UPR), an essential stress response needed during increased protein production (29, 30). While canonically considered to be induced in newly formed ASC, recent work indicates that activated B cells (actB) upregulate an array of UPR-affiliated genes. This process is controlled by mTORC1 prior to antibody production and before XBP1 activity (31), a known regulator of UPR in ASC (32, 33). Moreover, single-cell RNA-sequencing (scRNA-seq) of actB uncovered an IRF4-dependent bifurcation event that committed a portion of actB to ASC during the early stages of B cell activation (17). Thus, while recent work has highlighted critical early reprogramming events in actB, the timing and extent to which the above factors, and others, remains to be fully understood and integrated.

In this study, we aimed to understand the IRF4-dependent division-coupled reprogramming events that occur during the initial stages of B cell differentiation. Using an in vivo model of B cell differentiation (25), we found that IRF4-deficient B cells begin to divide normally but stall during the proliferative response. To assess the timing and scope of IRF4-dependent reprogramming, IRF4-sufficient and -deficient B cells at discrete divisions were sorted for RNA-seq and the assay for transposase accessible chromatin-sequencing (ATAC-seq) (34, 35). RNA-sequencing revealed that early upregulation of gene sets critical for ASC formation were dependent on IRF4. These included MYC target genes and genes important for OXPHOS. Indeed, IRF4-deficient B cells failed to fully upregulate *Myc* and displayed altered cell cycle distribution. The activity of mTORC1 was also reduced, resulting in an inability of IRF4-deficient B cells to undergo cell growth and initiate the UPR (31). ATAC-seq identified hundreds of differentially accessible regions (DAR) and established a hierarchy of IRF4 activity, with AP-1:IRF (AICE motifs) active during early divisions and ETS:IRF (EICE) motifs active in later divisions. Together, these data create a road map defining the role of IRF4 during the earliest stages of B cell differentiation in vivo and reveal a critical role for IRF4 in controlling cell growth and maintaining the proliferative response.

## Materials and Methods

### Mice and adoptive transfers

*Cd19*^Cre^ (JAX; 006785)(36) and *Irf4*^fl/fl^ (JAX; 000664)(16) mice were purchased from The Jackson Laboratory and bred to generate *Cd19*^Cre/+^*Irf4*^fl/fl^. CD45.2 μMT (JAX; 008100)(37) were bred onto the CD45.1 background to obtain CD45.1 μMT mice (17). All experimental animals were between 7 - 12 weeks of age and genders were equally represented. For adoptive transfers, naïve splenic CD43^−^ B cells were magnetically isolated using the B cell isolation kit (Miltenyi Biotec, Inc.; 130-090-862) and LS columns (Miltenyi Biotec, Inc.; 130-042-40). Isolated B cells were stained with CellTrace Violet (CTV) (Life Technologies; C34557) per the manufacturer’s protocol and resuspended in sterile PBS (Corning; 21-040-CV) before transferring 15×10^6^ B cells into a disparate congenic μMT host. At 24 h post-transfer, host mice were challenged intravenously with 50 μg LPS (Enzo Life Sciences; ALX-581-008), intranasally with 0.1 LD_50_ influenza A/HK-X31 (X31), or intravenously with 50 μg NP-Ficoll (Biosearch Technologies; F-1420-10). For influenza infections, mice were anesthetized with vaporized isoflurane (Patterson Veterinary; 07-893-1389) before X31 administration. Experimental mice were euthanized via carbon dioxide asphyxiation in accordance with AVMA guidelines. All procedures were approved by the Emory Institutional Animal Care and Use Committee.

### Flow cytometry and sorting

Cells were resuspended at 1×10^6^ / 100 μl in FACS buffer (1X PBS, 1% BSA, and 2 mM EDTA), stained with Fc Block (BD; 553141) and antibody-fluorophore conjugates for 15 and 30 m, respectively, and then washed with 1 ml of FACS. For adoptive transfers when NP-Ficoll or X31 was used, CD45.2 transferred cells were enriched prior to antibody staining using anti-CD45.2-APC or anti-CD45.2-PE followed by magnetic enrichment using anti-APC (Miltenyi; 130-090-855) or anti-PE (Miltenyi; 130-097-054) microbeads. The following antibody-fluorophore conjugates and stains were used: B220-PE-Cy7 (Biolegend; 103222), B220-A700 (Biolegend; 103232), BrdU-APC (Biolegend; 339808), c-MYC-PE (Cell Signaling; 14819), c-MYC-Alexa Fluor 647 (Cell Signaling; 13871), CD11b-APC-Cy7 (Biolegend; 101226), CD138-BV711 (BD; 563193), CD138-APC (Biolegend; 558626), CD45.1-FITC (Tonbo Biosciences; 35-0453-U500), CD45.1-PE (Biolegend; 110708), CD45.1-APC (Biolegend; 110714), CD45.1-APC-Cy7 (Tonbo Biosciences; 25-0453-U100), CD45.2-PE-Cy7 (Biolegend; 109830), CD45.2-PerCP-Cy5.5 (Tonbo Biosciences; 65-0454-U100), CD45.2-PE (Tonbo Biosciences; 50-0454-U100), CD45.2-APC (Biolegend; 109814), CD90.2-APC-Cy7 (Biolegend; 105328), F4/80-APC-Cy7 (Biolegend; 123118), Fas-PerCP-Cy5.5 (Biolegend; 152610), GL7-eFluor 660 (Fisher Scientific; 50-112-9500), GL7-PerCP-Cy5.5 (Biolegend; 144610), GL7-PE-Cy7 (Biolegend; 144620), Ki67-APC (Biolegend; 652406), pS6-PE (Cell Signaling; 5316), Rabbit mAb IgG XP Isotype-Alexa Fluor 647 (Cell Signaling; 2985), Rabbit mAb IgG XP Isotype-PE (Cell Signaling; 5742), Zombie Yellow Fixable Viability Kit (Biolegend; 423104), Zombie NIR Fixable Viability Kit (Biolegend; 423106), CellTrace Violet (Life Technologies; C34557), and 7AAD (Biolegend; 76332). For all flow cytometry analyses involving adoptive transfers, the following general gating strategy was used: lymphocytes were gated based on SSC-A / FSC-A, single cells by FSC-H / FSC-W or FSC-H / FSC-A, live cells based on exclusion of Zombie Yellow or Zombie NIR Fixable Viability Kit, and the markers CD11b, F4/80, and CD90.2 to remove non-B cells. All flow cytometry were performed on an LSR II, LSRFortessa, or LSR FACSymphony (BD) and analyzed using FlowJo v9.9.5, v10.5.3, or v10.6.2. Cell sorting was performed at the Emory Flow Cytometry Core using a FACSAria II (BD) and BD FACSDiva software v8.0.

### Cell cycle analysis and intracellular staining

In some adoptive transfers, hosts were injected with 800 μg BrdU (Biolegend; 423401) intravenously 1 h prior to euthanasia. Staining of BrdU, Ki67, and 7AAD was achieved using the Phase-Flow BrdU Cell Proliferation Kit (Biolegend; 370704), substituting anti-BrdU for anti-Ki67 when desired. Intracellular pS6 staining was accomplished following BD’s two-step protocol using BD Phosflow Fix Buffer I (BD; 557870) and BD Phosflow Perm Buffer III (BD; 558050). As a negative control for intracellular pS6, cultured cells were treated with 200 nM of rapamycin (Sigma-Aldrich; R8781) for 2 h prior to staining. Intracellular staining of MYC was performed using the FIX & PERM Cell Permeabilization Kit (ThermoFisher; GAS003) per the manufacturer’s protocol.

### Ex vivo B cell differentiation

Isolated B cells were cultured at a concentration of 0.5 × 10^6^ cells/ml in B cell media (RPMI 1640 supplemented with 1X nonessential amino acids, 1X penicillin/streptomycin, 10 mM HEPES, 1 mM sodium pyruvate, 10% heat-inactivated FBS, and 0.05 mM 2-ME) containing 20 mg/ml *Escherichia coli* O111:B4 derived LPS (Sigma-Aldrich; L2630), 5 ng/ml IL-5 (Biolegend; 581504), and 20 ng/ml IL-2 (Biolegend; 575406) as previously described (38). Additional LPS (10 μg/ml), IL-5 (2.5 ng/ml), and IL-2 (10 ng/ml) were added to the cultures every 24 h for the duration of the time course.

### Retroviral production and transduction

Retrovirus was prepared as previously described (39). Briefly, Platinum-E cells were transfected at 70-80% confluency on 10 cm plates with 4 μg pCL-Eco(40) and 6 μg of either pMSCV-pBabeMCS-IRES-RFP (Addgene; 33337) or pMSCV-Myc-IRES-RFP (Addgene; 35395)(41) using 40 μl TransIT-293 (Mirus; MIR2700). Cell media (antibiotic-free DMEM supplemented with 10% heat-inactivated FBS) was replaced with High-BSA cell media (DMEM supplemented with 10% heat-inactivated FBS and 1g/100ml BSA) 18 h after transfection. Retrovirus was harvested 24 and 48 h later, filtered through a 0.45 μm membrane, and concentrated using 5x PEG-it viral precipitation solution (System Biosciences; LV825A-1). Transduction of B cells was performed 12-24 h after stimulation via spinfection at 800 g for 1 h.

### Quantitative RT-PCR

One million cells were resuspended in 600 μl of RLT Buffer (Qiagen; 79216) containing 1% 2-BME and snap frozen in a dry ice – ethanol bath for RNA isolation. Lysates were thawed, subjected to QIAshredder homogenization (Qiagen; 79656), and then total RNA isolation using the RNeasy Mini Ki (Qiagen; 74104). RNA was reverse transcribed using SuperScript II Reverse Transcriptase (Invitrogen; 18064014). cDNA was diluted 1 μg / 100 ul and qPCR was performed on a CFX96 Instrument (Bio-Rad) using SYBR Green incorporation. Primers used included: 18S-forward 5’-GTAACCCGTTGAACCCCATT-3’ 18S-reverse 5’-CCATCCAATCGGTAGTAGCCG-3’, MYC-forward 5’-CGATTCCACGGCCTTCTC-3’, and MYC-reverse 5’-TCTTCCTCATCTTCTTGCTCTTC-3’. All primers were purchased from Integrated DNA Technologies.

### RNA-sequencing and data analysis

For all samples, 1,000 cells were sorted into 300 μl of RLT buffer (Qiagen; 79216) containing 1% 2-ME and snap frozen in a dry ice – ethanol bath. RNA isolation was achieved using the Quick-RNA Microprep kit (Zymo Research; R1050). Isolated RNA was used as input for the SMART-seq v4 cDNA synthesis kit (Takara; 634894), and 400 pg of cDNA was used as input for the NexteraXT kit (Illumina). Final libraries were quantified by qPCR and bioanalyzer traces, pooled at equimolar ratios, and sequenced at the New York University Genome Technology Center on a HiSeq 4000.

Raw sequencing data were mapped to the mm10 genome using STAR v.2.5.3 (42). Duplicate reads were identified and removed using PICARD (http://broadinstitute.github.io/picard/). The Bioconductor package edgeR v3.24.3 (43) was employed to determine differentially expressed genes (DEG), which were defined as having an absolute log_2_ fold-change of ≥1 and a false discovery rate (FDR) of ≤0.05. All detected transcripts were ranked by multiplying the sign of fold change (+/–) by -log_10_ of the p-value, and gene set enrichment analysis (GSEA) (44) was performed on this ranked gene list. All t-SNE projections were generated using ‘Rtsne’ v 0.15 (https://github.com/jkrijthe/Rtsne). Clustering and heatmap analysis were achieved using ‘heatmap3’ (https://github.com/cdschar/heatmap).

### ATAC-sequencing and data analysis

For each sample, 10,000 cells were sorted into FACS buffer and the assay for transposase-accessible chromatin sequencing (ATAC-seq) was performed. Tn5 preparation and library generation was previously described (23). Briefly, cells were centrifuged at 500 g for 10 min at 4 °C. The supernatant was removed and cells were resuspended in 25 μl of Tn5 tagmentation reaction (2.5 μl Tn5, 12.5 μl 2X tagmentation buffer (20 mM TAPS-NaOH pH 8.1, 10 mM MgCl2, 20% DMF), 2.5 μl 1% Tween-20, 2.5 μl 0.2% digitonin, and 5 μl of molecular grade water). Resuspended samples were incubated at 37°C for 1 h. Cells were then lysed by adding 25 μl lysis buffer (300 mM NaCl, 100 mM EDTA, 0.6% SDS, and 2 μl 10 mg/ml proteinase K) and incubated for 30 min at 40°C. Transposed DNA was isolated using AMPure XP SPRI beads (A63880) by adding 0.7x volumes to remove high molecular weight DNA and then 1.2x volumes to positively select for low molecular weight DNA. Tagmented DNA was eluted in 15 μl EB buffer (Qiagen; 19086) and amplified using Nextera indexing primers (Illumina) and KAPA HiFi polymerase (Roche; KK2601). Final libraries were sequenced at the New York University Genome Technology Center on a HiSeq 4000.

Raw sequencing data were mapped to the mm10 genome using Bowtie v1.1.1 (45). Peaks were called using MAC2 v 2.1.0 (46) and annotated to the nearest gene using HOMER v4.8.2 (47). Reads per peak million normalization was performed for all samples as previously described (35). The Bioconductor package edgeR v3.24.3 (43) was used to determine differentially accessible regions (DAR), which were defined as having an absolute log_2_ fold-change of ≥1 and a FDR of ≤ 0.05. Motif analysis was performed using the HOMER program findMotifsGenome.pl (de novo results). For plotting the rank value of transcription factors, enriched transcription factor motifs were ranked according to their p-value and normalized by the total number of enriched motifs found for a given sample. Resulting values were z-score normalized and motifs binned according to their DNA binding domain family.

### Statistics

All statistical analyses were achieved by using R/Bioconductor v3.5.2, Microsoft Excel v16.36 or v16.48, and GraphPad Prism v6.0c, v8.4.1, or 8.4.3. P values of less than 0.05 were considered significant. For RNA- and ATAC-seq significance, a combination of FDR and fold-change was used to designate DEG and DAR.

### Data availability

All sequencing data generated in this study have been deposited in NCBI Gene Expression Omnibus (https://www.ncbi.nlm.nih.gov/geo/) under accession code GSE173437 (GSE173435 for ATAC-seq and GSE173436 for RNA-seq).

## Results

### IRF4-deficient B cells responding to LPS in vivo stall during the proliferative response

Cell division is one of the earliest events following B cell activation, however a complete understanding of factors that control or maintain the proliferative response remain to be determined. Recent work identified an IRF4-dependent bifurcation event in the earliest stages of B cell activation (17). Cells along the IRF4-dependent branch upregulated gene sets critical for proliferation, indicating IRF4 may be important for controlling the proliferative response in vivo. To explore if IRF4 impacted cell proliferation during B cell differentiation, an in vivo adoptive transfer model was applied (25). Here, splenic naïve B cells from CD45.2^+^*Cd19*^+/+^*Irf4*^fl/fl^ (Ctrl) or CD45.2^+^*Cd19*^Cre/+^*Irf4*^fl/fl^ (IRF4cKO) mice were isolated, labeled with CellTrace Violet (CTV), and transferred to CD45.1^+^ μMT hosts. After 1 day, host mice were challenged with the type I T cell independent antigen LPS and cell division and differentiation were determined via CD138 expression (48, 49) in a time course covering three days (**Fig. 1A**). At 24 h, no division was observed for Ctrl and IRF4cKO cells, indicating a similar delay before initiating the proliferative response (**Fig. 1B**). At 48 h, both Ctrl and IRF4cKO cells began to divide, and the majority of responding cells were observed in divisions 2-4. A modest difference in IRF4cKO B cells in divisions 0-1 was observed at this time point (**Fig. 1B, 1C**). At 60 h, Ctrl were distributed in all cell divisions (0-8), with a subset differentiating after reaching or exceeding division 8. Comparatively, IRF4cKO cells accumulated in divisions 2-4, with few cells observed in divisions 5 and 6 (**Fig. 1B, 1C**). Strikingly, while more than half of Ctrl cells accumulated in division 8 at 72 h, the cell division pattern for cells from IRF4cKO largely remained the same as their 60 h time point, indicating the IRF4cKO cells stalled during the proliferative response (**Fig. 1B, 1C**). Indeed, the mean division number (MDN) (50) for Ctrl cells increased by ∼2 divisions from 60 to 72 h, while the MDN for IRF4cKO cells was unchanged (**Fig. 1D**). This proliferative defect was also reflected in reduced frequency of IRF4cKO cells detected in host spleens at 72 h (**Fig. 1E, 1F**). Importantly, staining for the pro-apoptotic marker annexin V revealed no differences in apoptosis or necrosis at 72 h in vivo (**Supplemental Fig. 1**). Furthermore, no differences in homeostatic proliferation were observed in mice that received Ctrl or IRF4cKO B cells but no LPS (**Fig. 1B**). It is also important to note that the vast majority of the splenic cells transferred divided at least once to LPS stimulation, indicating that nearly all B cells and not just a subset were responding in vivo. Proliferation defects were also observed when C57BL/6J mice were used as hosts (**Supplemental Fig. 2**). These data indicate IRF4 controls the proliferative capacity of B cells in response to LPS immune challenge.

**FIGURE 1.**
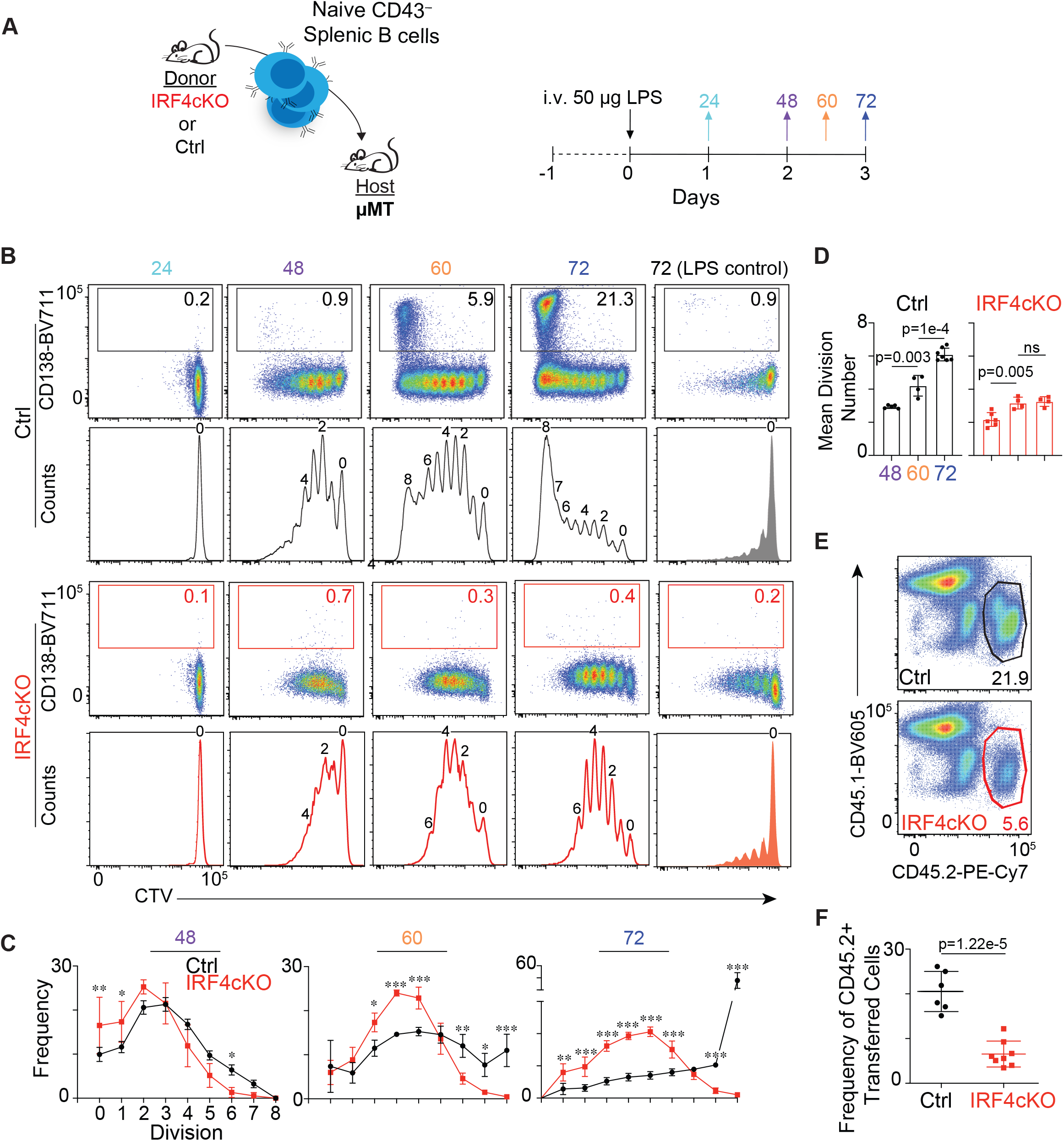
IRF4-deficient B cells stall during the proliferative response to LPS. **(A**) Schematic of experimental design. Ctrl (CD45.2^+^*Cd19*^+/+^*Irf4*^fl/fl^) or IRF4cKO (CD45.2^+^*Cd19*^Cre/+^*Irf4*^fl/fl^) splenic B cells were CTV-labeled and adoptively transferred into µMT (CD45.1+) mice, as described in the methods. At 24 h post transfer, mice were inoculated with LPS i.v. At the indicated time points, spleens were harvested and analyzed. (**B**) Flow cytometry histograms displaying cell division and ASC differentiation (CD138^+^). The frequency of CD138+ cells are shown. **(C)** Frequency of transferred (CD45.2^+^) cells at discrete divisions for 48, 60, and 72 h. **(D)** Mean division number of all responding cells at each time point. (**E)** Ctrl (top) and IRF4cKO (bottom) representative flow cytometry plots of CD45.1 versus CD45.2 with gates drawn and frequencies shown for the transferred population. **(F)** Quantification of the frequency of CD45.2 transferred cells from **E**. All data are representative of at least two independent experiments using at least 3 mice per group. Data in **C, D**, and **F** represent mean ± SD. Statistical significance in **C** was determined by a two-way ANOVA with Sidak’s multiple comparisons test. Statistical significance in **D** was determined by a paired two-tailed Student’s *t* test, while statistical significance in **F** was determined by determined by a two-tailed Student’s *t* test. * p < 0.05, ** p < 0.01, *** p <0.001.

**FIGURE 2.**
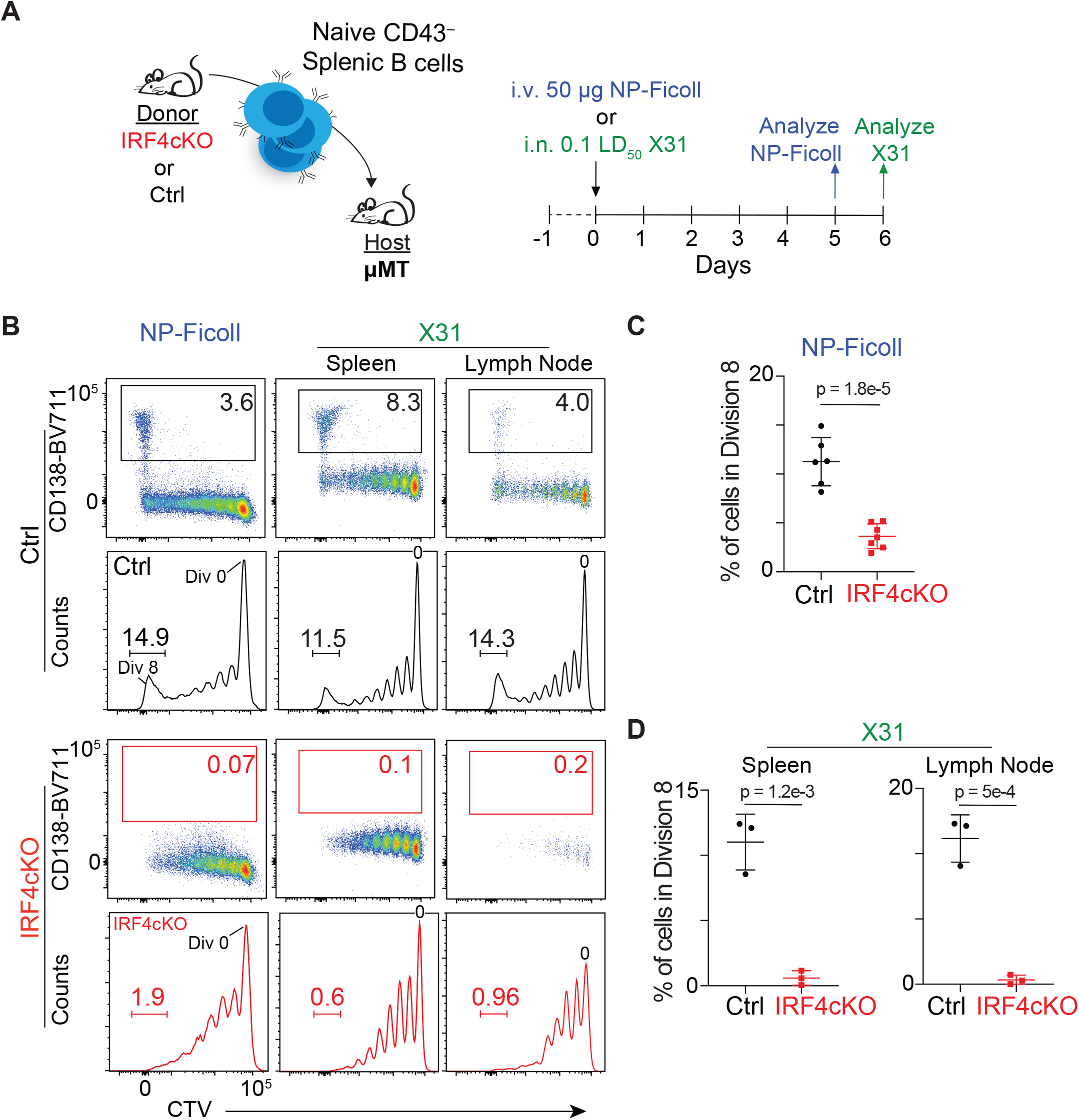
IRF4-deficient B cells exhibit a proliferation defect in response to T-independent and T-dependent antigens. (**A**) Schematic of experimental design. Ctrl and IRF4cKO B cells were prepared and adoptively transferred as in Fig 1 and the methods section. Here, animals were stimulated with either NP-Ficoll or infected with influenza strain X31 as described in the methods. Spleens from NP-Ficoll inoculated animals were harvested at d5; and for influenza, both spleens and the draining mediastinal lymph nodes were isolated at d6 post-challenge. **(B)** Representative flow cytometry plots of CD138 versus CTV or CTV histograms for Ctrl and IRF4cKO. The frequency of CD138^+^ (top) and division 8 (bottom) cells are shown. Frequency of division 8 cells for Ctrl and IRF4cKO from **B** following NP-Ficoll (**C**) or influenza X31 (**D**) challenge. All data are representative of two independent experiments using at least 3 mice per group. Data in **C** and **D** represent mean ± SD with statistical significance determined by a two-tailed Student’s *t* test.

### IRF4-deficient B cells exhibit a proliferation defect to T-independent and -dependent antigens

To determine whether IRF4 controls the proliferative response to other stimuli, adoptive transfers were performed followed by challenge with the type II T-independent antigen 4-hydroxy-3-nitrophenylacetyl (NP)-Ficoll or the T-dependent antigen influenza A/HK-X31 (X31). Five days post-NP-Ficoll and six day after X31 challenge, host mice were sacrificed, and cell division and differentiation were assessed by flow cytometry (**Fig. 2A**). Because NP-Ficoll and X31 stimulate antigen-specific B cells that represent a small portion of the population, the majority of Ctrl and IRF4cKO cells remained undivided for both stimulation conditions (**Fig. 2B**). For NP-Ficoll, Ctrl cells were distributed in all cell divisions 1-8, and a subset of cells that reached or surpassed division 8 differentiated (**Fig. 2B, 2C**). Similar results were observed following X31 challenge and independent of whether the transferred cells were recovered in the mediastinal lymph node or the spleen (**Fig. 2B, 2D**). Interestingly, CD138+ ASC were observed at division eight for all three antigen conditions for Ctrl cells. Comparatively, cells from IRF4cKO were mainly distributed in the first few divisions for both stimulation conditions, with very few IRF4cKO B cells detected after division 4 and almost none reaching division 8 and forming ASC (**Fig. 2B-D**). Taken together, these data indicate IRF4 plays a critical role in controlling the proliferative response to type II T independent and early T dependent antigen responses.

### IRF4-deficient B cells display altered cell cycle distribution

To better understand the proliferative defect observed above, the role that IRF4 played with respect to cell cycle was investigated. CTV-labeled Ctrl and IRF4-deficient B cells were adoptively transferred into µMT mice and recovered 72 h post-LPS stimulation. Cells were stained with Ki-67 and 7AAD to distinguish the frequency of cells in each phase of the cell cycle at discrete divisions (51) and analyzed by flow cytometry. These data revealed that in the final detectable divisions, IRF4cKO cells accumulated in G_0_/G_1_ with a corresponding decrease in cells found in the G_2_/M (**Fig. 3A, 3B**). This was in stark contrast to Ctrl cells, which revealed more cells in S and G2/M at the same divisions. This indicates that the cell cycle was significantly perturbed in B cells from IRF4cKO in these final divisions (**Fig. 3A, 3B**). To better understand the proliferative defect observed in IRF4cKO cells in vivo, the frequency of actively proliferating cells by BrdU incorporation was examined after IRF4cKO cells had stalled. Appreciably, a lower frequency of BrdU^+^ IRF4cKO compared to Ctrl cells were observed (**Fig. 3C, 3D**). BrdU+ IRF4cKO cells were also distributed proportionally to the total population. In contrast, BrdU+ Ctrl cells were largely distributed in division 8 (**Fig. 3C**). Thus, IRF4 is critical for cell cycle control and maintaining the proliferative response.

**FIGURE 3.**
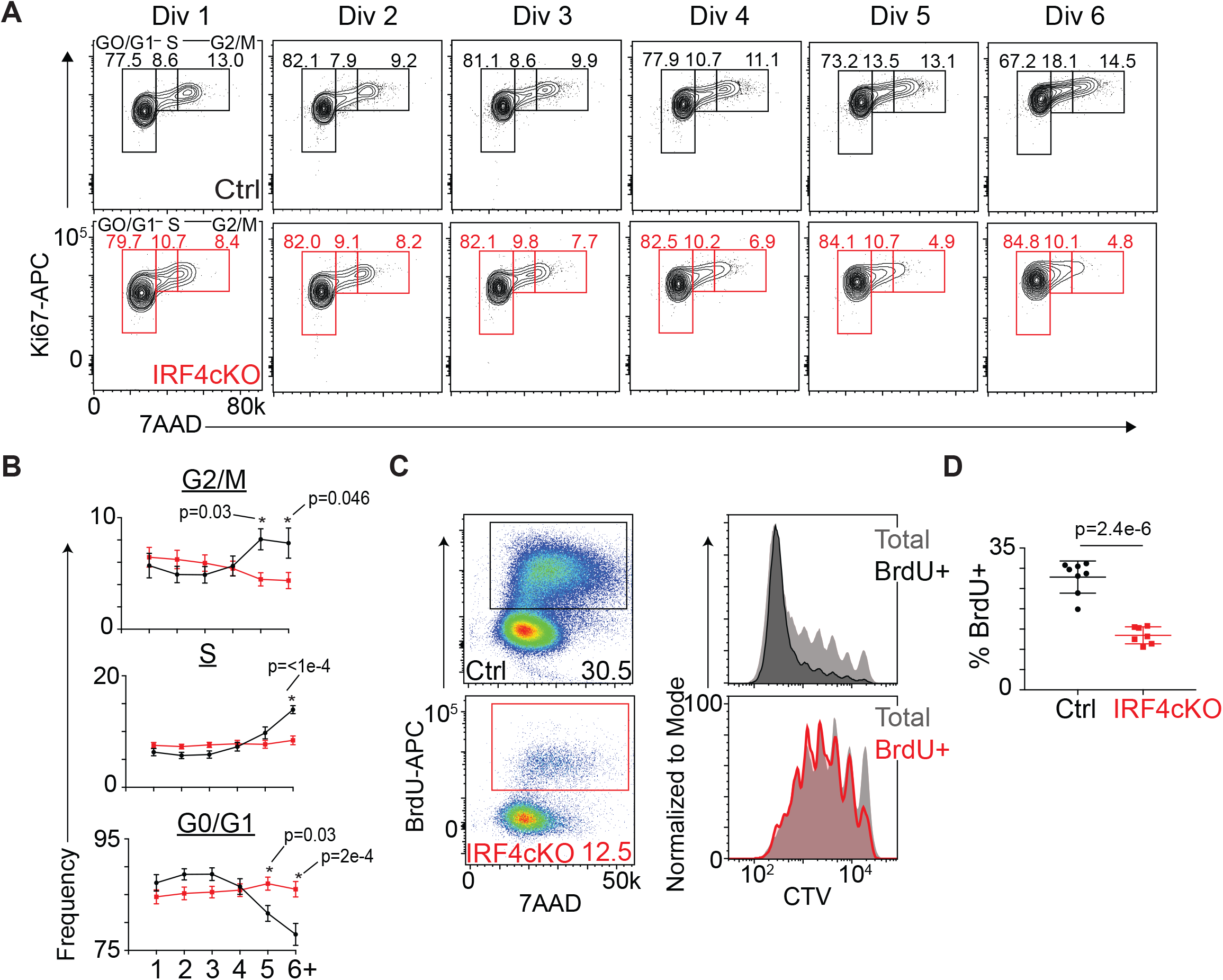
IRF4-deficient B cells display altered cell cycle distribution. **(A)** Ctrl (black) and IRF4cKO (red) B cells were prepared, adoptively transferred, and inoculated with LPS as in Fig 1. At 72 h, mice were sacrificed and the spleens harvested. Cells were stained with Ki67 and 7AAD and representative flow cytometry plots at the indicated divisions are shown. Flow cytometry gates indicating G0/G1, S, and G2/M phase of the cell cycle are shown with the frequency of cells for each. **(B)** Quantification of the data from **A** displaying the frequency of cells found in each phase of the cell cycle at each division. **(C)** Following the above adoptive transfer scheme described in **A**, mice were injected with BrdU 1 h prior to sacrifice to assess active S phase of the cell cycle. Representative flow cytometry plot of BrdU versus 7AAD (left) and CTV histograms (right) of the total transferred population (grey) overlaid with the BrdU^+^ cells to visualize the distribution of actively proliferating cells. **(D)** Quantification of the data from **C** displaying the frequency of BrdU^+^ cells. All data are representative of at least two independent experiments using at least 3 mice per genotype. Data in **B** and **D** represent mean ± SD. Statistical significance in **D** was determined by a two-tailed Student’s *t* test. Statistical significance in **B** was determined by a two-way ANOVA with Sidak’s multiple comparisons test. P-values are shown at points of significance.

### Cell division-coupled IRF4-dependent transcriptional reprogramming

B cell differentiation to ASC requires considerable transcriptional rewiring that consists of progressive cell division-based reprogramming events (25). To determine the impact of IRF4 on this process, Ctrl and IRF4cKO cells were sorted from divisions 0, 1, 3, 4, 5, and 6 as determined by CTV dilution (**Fig. 4A**) and subjected to RNA-seq analyses. Comparing gene expression profiles for Ctrl and IRF4cKO cells in the same division revealed hundreds of differentially expressed genes (DEG) that increased or decreased expression in IRF4-deficient B cells, indicating IRF4 functions to repress and activate gene expression programs, even in the earliest stages of actB reprogramming (**Fig. 4B**). This activity is consistent with previous work, demonstrating that a significant increase in IRF4 levels occurrs after the first cell division (17, 18). After successive divisions, IRF4cKO B cells became progressively transcriptionally divergent compared to Ctrl cells (**Fig. 4B**). Hierarchical clustering of samples reflected this divergency with Ctrl and IRF4cKO samples in divisions 0 and 1 clustering by gene expression and divisions 3 - 6 clustering by IRF4 status (**Fig. 4C**). T-distributed stochastic neighbor-embedded (t-SNE) projections of gene expression data from all samples indicated major cell division-dependent transcriptional reprogramming events that were dependent on IRF4 and predominately in divisions 3 - 6. (**Fig. 4D**). Collectively, IRF4cKO are transcriptionally distinct by division 3 and continue to diverge through subsequent divisions. Thus, cell division-based IRF4-dependent reprogramming occurs during the initial stages of B cell differentiation.

**FIGURE 4.**
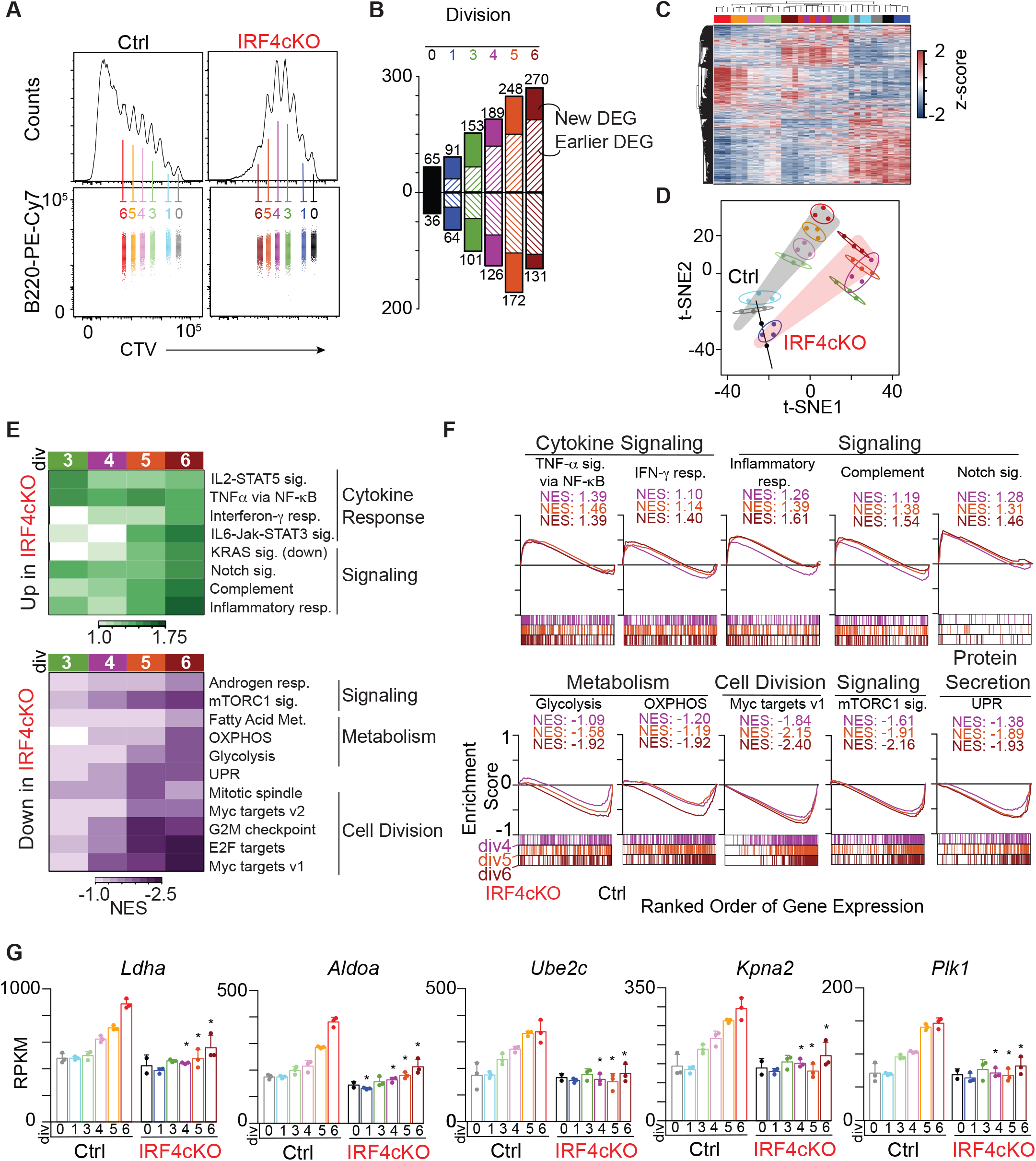
IRF4-deficient B cells fail to upregulate metabolic and proliferative gene expression programs during B cell differentiation. **(A)** Ctrl and IRF4cKO B cells were prepared, adoptively transferred, and inoculated with LPS as in Fig 1 and harvested at 72 h. Cells at the indicated divisions were sorted and subjected to RNA-seq as described in methods. Representative flow cytometry plots of B220 and CTV histograms and projections of the sorted populations are shown and labeled by division number. **(B)** Bar plot quantifying the number of differentially expressed genes (DEG) at each division that increase (top) or decrease (bottom) expression in IRF4cKO cells compared to Ctrl. Solid bars indicate the proportion of genes that represent a new DEG appearing in that division while striped bars indicate the proportion of genes that were a DEG in an earlier division. **(C)** Hierarchical clustering of the expression of 10,404 genes detected from **A. (D)** t-SNE projections of RNA-seq data from control samples (highlighted in grey) and IRF4cKO samples (highlighted in red). **(E)** Heat map of normalized enrichment scores (NES) calculated by gene set enrichment analysis (GSEA) (44) for pathways upregulated and downregulated in IRF4cKO. **(F)** GSEA examples for the indicated gene sets for IRF4cKO up and down DEG from divisions 4, 5, and 6. NES values are indicated for each division. **(G)** Bar plot displaying reads per kilobase million (RPKM) values for the indicated genes at all sequenced divisions for Ctrl and IRF4cKO cells. Asterisks above IRF4cKO division data indicate significance (FDR < 0.001) when compared to the corresponding Ctrl division. Data were derived from 3 independent adoptive transfers for Ctrl and IRF4cKO. One division 0 IRF4cKO sample was excluded due to a high frequency of duplicate reads.

To determine the transcriptional programs dependent on IRF4, gene set enrichment analysis (GSEA) (44) was performed for DEG that increased or decreased expression in IRF4cKO cells in divisions 3 - 6. IRF4cKO B cells progressively failed to induce gene sets important for cell division, metabolism, and signaling (**Fig. 4E, 4F**). This consisted of genes critical for glycolysis and OXPHOS, which are critical metabolic programs for actB and ASC, respectively (24, 52) (**Fig. 4E, 4F**). Enzymes that failed to be induced and are critical for glycolytic metabolism included *Ldha* (53) and *Aldoa* (54) (**Fig. 4G**). Additionally, mTORC1 signaling and MYC target genes failed to be induced in IRF4cKO cells, and included genes that promote cell proliferation such as *Ube2c* (55), *Kpna2* (56), and *Plk1* (57, 58) (**Fig. 4G**). Notably, the cell cycle was significantly perturbed in IRF4cKO cells in the divisions in which MYC target genes were the most dysregulated (**Fig. 3A, 3B**). These data are consistent with reports that reduction of *Myc* impacts G1-S transition of the cell cycle (59-61). Genes sets that failed to be repressed consisted of those involved in cytokine and cell signaling, such as the inflammatory response, and reflect previous reports that IRF4-deficient B cells progress down a reprogramming path whose gene expression program reflects cells responding to inflammatory stimuli (17). Collectively, these data suggest that early metabolic and proliferative programs essential for cell growth and division are dependent on IRF4.

### ATAC-sequencing reveals a hierarchy of IRF4 activity

To identify regions that change chromatin accessibility during B cell differentiation upon deletion of *Irf4*, paired ATAC-seq (62) data derived from the above divisions was analyzed to reveal IRF4-specific regulatory activities and IRF4-dependent transcription factor networks that impact B cell differentiation. Comparison of Ctrl and IRF4cKO cells in discrete divisions identified hundreds of differentially accessible regions (DAR), with a progressive increase in DAR occurring after the first cell division and more than 700 DAR by divisions 5 and 6 (**Fig. 5A**). These differences were also reflected in t-SNE spatial projections (**Fig. 5B**), and indicated that similar to RNA-seq, chromatin accessibility differences occurred predominately in divisions 3 - 6 (**Fig. 5A, 5B**). Collectively, these data support the notion that IRF4-dependent reprogramming occurs progressively beginning during the initial stages of B cell differentiation and that the chromatin landscape of IRF4cKO B cells is markedly distinct by division 3.

**FIGURE 5.**
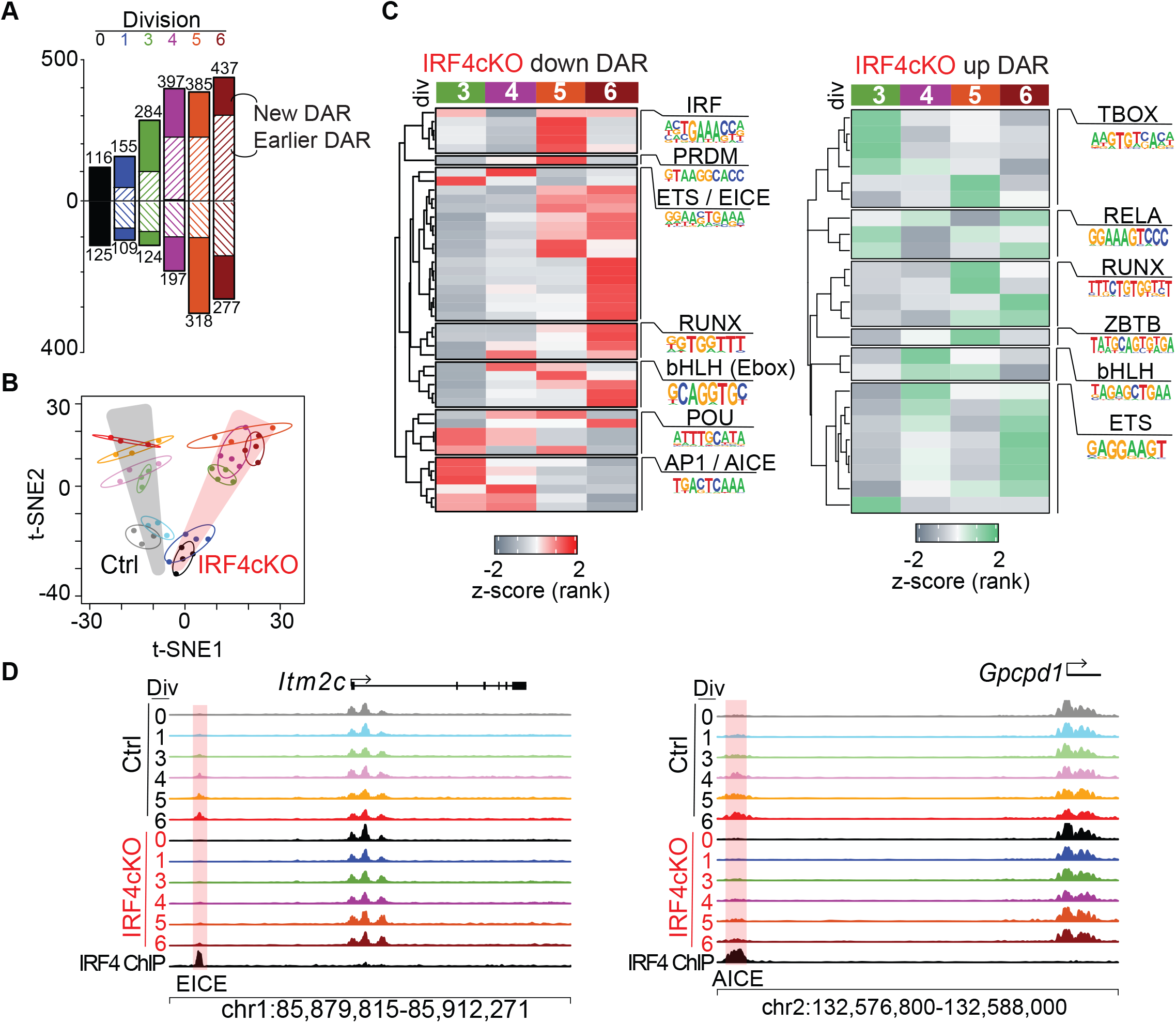
IRF4-deficient B cells display progressively altered chromatin accessibility profiles after subsequent divisions. ATAC-seq was performed on the sorted cell populations described in Figure 4. **(A)** Bar plot quantifying the number of differentially accessible regions (DAR) at each division that increase or decrease in IRF4cKO compared to Ctrl. Solid bars indicate the proportion of DAR that are new to that division, while striped bars indicate the proportion of regions that were a DAR in an earlier division. **(B)** t-SNE plots of 8,005 accessible loci from Ctrl samples (highlighted in grey) and IRF4cKO samples (highlighted in red). **(C)** Heatmap of HOMER (47) rank scores (by division) for the top 10 transcription factor motifs and related family members identified in IRF4cKO division 6 DAR. TF family names and a representative motif are displayed in their respective group. **(D)** ATAC accessibility profile for the indicated regions at DAR with an EICE (left) and AICE (right) motif. DAR regions are highlighted in red. IRF4 ChIP-seq from Minnich et al (93) was included in the IRF4 track. ATAC-seq data were derived from 3 independent adoptive transfers for Ctrl and 4 independent adoptive transfer for IRF4cKO. One division 5 IRF4cKO sample was excluded due to low coverage.

To gain a better understanding of the transcription factor networks dependent on IRF4, the top 10 enriched DNA sequence motifs in division 6 DAR were determined and matched to known putative transcription factor binding motifs using HOMER (47). Because enrichment p-values are dependent on the number of DAR, each transcription factor motif was rank normalized based on significance at each division, and the change in rank score across the divisions plotted, revealing how motif accessibility was altered across the divisions. Motifs enriched in regions that decreased accessibility in IRF4cKO cells (down DAR) included known IRF4 DNA binding motifs (14, 15, 18, 63), such as the core IRF motif (GAAA), AP-1-IRF composite element (AICE) (64), and ETS-IRF composite element (EICE) (65, 66) (**Fig. 5C**). Interestingly, this revealed a hierarchy among heterodimeric IRF4 binding sites (67), with AICE more highly ranked in early divisions and EICE motifs more highly ranked in later divisions. DAR in proximity of *Itm2c* and *Gpcpd1* reflected this hierarchy of activity (**Fig. 5D**). These data support the kinetic control of IRF4 activity (18, 27), as well as previous work implicating the timing of IRF4 in conjunction with the AP-1 transcription factor BATF in early cell fate decisions during B cell differentiation (17). Other transcription factors enriched in down DAR in the final divisions included RUNX and E-box binding bHLH family members (**Fig. 5C**).

Among regions that increased accessibility in IRF4cKO (up DAR), TBOX family members were more highly ranked in early divisions compared to subsequent divisions (**Fig. 5C**). Notably, the TBOX family member TBET supports ASC formation through repression of the inflammatory gene expression program (68), which was progressively upregulated in IRF4cKO cells (**Fig. 4E, 4F**). RUNX and ETS family members were most highly ranked in the final divisions, suggesting that these transcription factors are playing roles at both regions gaining and losing accessibility as the cells differentiate (**Fig. 5C**). Collectively, these data demonstrate the timing of IRF4-dependent reprogramming, establish a hierarchy of IRF4 activity that occurs at early and late cell divisions, and identify transcription factor networks dependent on IRF4.

### IRF4-deficient B cells fail to upregulate MYC

Recent work described MYC as a cell division timer during lymphocyte differentiation, with division cessation occurring when MYC levels fell below a critical threshold (12). We reasoned that *Myc* may be dysregulated in IRF4-deficient B cells because IRF4cKO cells: 1) stalled during the proliferative response to LPS (**Fig. 1**); 2) accumulated in G_0_/G_1_ phase of the cell cycle (**Fig. 3A, 3B**); 3) progressively failed to induce MYC target genes (**Fig. 4E, 4F**); and 4) E-box binding bHLH family members were enriched in down DAR in divisions where MYC target genes were the most dysregulated (**Fig. 5C**). In fact, IRF4cKO cells were progressively enriched for genes dysregulated in MYC-deficient B cells stimulated with LPS and IL4 (11), further supporting the notion that MYC programming is altered in IRF4cKO cells (**Fig. 6A**). To determine if *Myc* failed to be induced in IRF4-deficient B cells, Ctrl and IRF4cKO cells were cultured ex vivo with LPS, IL2, and IL5 to initiate the pathway to ASC (38), and expression was analyzed by RT-qPCR before and 24 h after stimulation. While no differences in *Myc* levels were detected prior to stimulation, a significant reduction was observed at 24 h (**Fig. 6B**). Similar observations were detected by intracellular staining of MYC, which confirmed that while MYC levels were increased over naïve B cells, IRF4cKO cells failed to upregulate MYC to the same level as Ctrl cells (**Fig. 6C, 6D**). These data are consistent with previous reports following PMA/IO treatment of IRF4-deficient and -sufficient B cells (69). The observed differences in MYC expression are likely caused by transcription of *Myc* and not due to alterations in MYC protein stability (70) (**Supplemental Fig. 3**).

**FIGURE 6.**
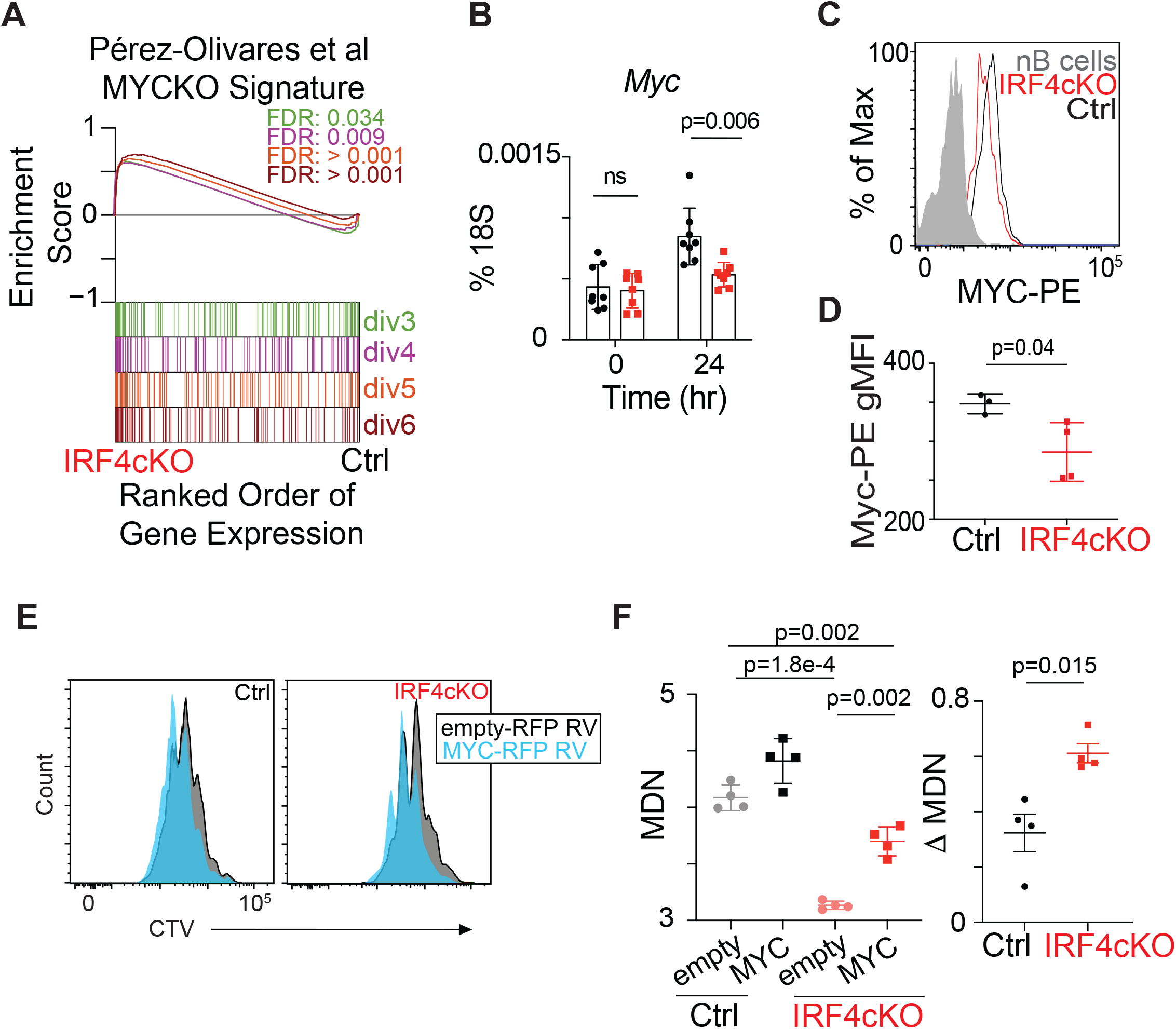
IRF4-deficient B cells fail to fully upregulate MYC. **(A)** GSEA using the top differentially expressed genes dysregulated in MYC-deficient B cells stimulated with LPS and IL-4 for 72 h (11). FDR values are displayed for each division as indicated by color. Splenic B cells from Ctrl and IRF4cKO mice were isolated and treated with LPS, IL2, and IL5 ex vivo as described in methods. **(B)** Quantitative RT-PCR expression of *Myc* relative to 18S rRNA expression before (0 h) or 24 h after stimulation. **(C)** Representative intracellular staining of MYC for naïve untreated B cells (nB) and 24 h stimulated Ctrl and IRF4cKO cells (top). (**D**) Geometric mean fluorescence intensity (gMFI) quantified for the stimulated samples for **C. (E)** Representative CTV histograms of Ctrl (left) and IRF4cKO (right) transduced with empty-RFP retrovirus (black) or MYC-RFP expressing retrovirus (blue). **(F)** (Left) Quantification of the mean division number (MDN) for Ctrl and IRF4cKO cells transduced with empty-RFP retrovirus or MYC-RFP retrovirus from **E**. (Right) Quantification of the change in MDN after MYC overexpression in Ctrl and IRF4cKO cells from **E**. All data are representative of at least two independent experiments using at least 3 mice per genotype. Data in **B, D**, and **F** represent mean ± SD. Statistical significance in **B** and **D** was determined by a two-tailed Student’s *t* test. Statistical significance in **F** when comparing IRF4cKO samples was determined by a paired two-tailed Student’s *t* test, while significance between Ctrl and IRF4cKO samples was calculated by a two-tailed Student’s *t* test.

To explore whether MYC overexpression could rescue the cell division defect of IRF4-deficient B cells, Ctrl and IRF4cKO cells were again cultured ex vivo with LPS, IL2, and IL5 and transduced with retrovirus expressing MYC-RFP or control RFP. Overexpression of *Myc* significantly improved the proliferation capacity of cells, and this improvement was greater for B cells from IRF4cKO than Ctrl (**Fig. 6E, 6F**). However, while IRF4cKO cells exhibited a greater proliferative gain upon MYC overexpression compared to Ctrl cells, full cell division capacity was not restored, as Ctrl B cells transduced with control RFP still displayed greater proliferative capacity. Collectively, these data suggest that IRF4cKO B cells fail to fine-tune the levels of *Myc* during the initial stages of B cell activation, which impact the overall cell division pattern and are consistent with the observation that IRF4cKO B cells begin to divide normally but stall in the middle of the proliferative response (**Fig. 1**). However, *Myc* overexpression alone does not fully restore the division capacity of IRF4cKO B cells, indicating additional deficiencies are contributing to the proliferative defect.

### IRF4-deficient B cells exhibit reduced mTORC1 activity and are unable to initiate the UPR

Activation of the mammalian target of rapamycin (mTOR) is essential for promoting biosynthetic processes necessary for cell growth and division (71). Importantly, ablation of mTORC1 activity impacted the proliferative effects of MYC overexpression in murine tumor cells (72), indicating there is significant crosstalk between the two signaling cascades (73-76). Recent work indicated mTORC1 coordinates an early B cell-activation unfolded protein response (UPR), in which a subset of UPR-affiliated genes are upregulated independent of XBP1 (31), a known driver of the UPR (33, 77). Interestingly, while Ctrl B cells gradually upregulated the B cell-activation UPR as early as division 3, IRF4cKO cells failed to initiate the program to the same levels (**Fig. 7A, 7B**). Indeed, genes associated with mTORC1 signaling progressively failed to be induced in IRF4cKO B cells (**Fig. 4E, 4F**). Collectively, these data implied that mTORC1 activation may be dysregulated in IRF4-deficient B cells. To test for mTORC1 activity, Ctrl and IRF4cKO cells were cultured ex vivo with LPS, IL2, and IL5 for 48 h, and intracellular staining for phosphorylation of the canonical mTORC1 substrate S6 (pS6) was performed. Strikingly, while the majority of B cells from Ctrl exhibited high amounts of pS6, most IRF4cKO cells contained pS6 levels similar to cultures where mTORC1 activity was blocked following treatment with rapamycin (**Fig. 7C, 7D**). Consistently, proliferating IRF4cKO cells also failed to increase in cell size compared to Ctrl B cells at 48 h post-LPS in vivo (**Fig. 7E, 7F**). Intriguingly, this reduction in cell size was rescued via overexpression of *Myc* in IRF4cKO cells cultured ex vivo (**Fig. 7G**). Thus, IRF4cKO B cells exhibit a defect in mTORC1 activity that impacts the ability of cells to increase in cell size that is overcome with *Myc* overexpression. Thus, these data support the role of mTORC1 in upregulating an early B cell-activation UPR, assign the cell division in which this process occurs, and implicate IRF4 in this process.

**FIGURE 7.**
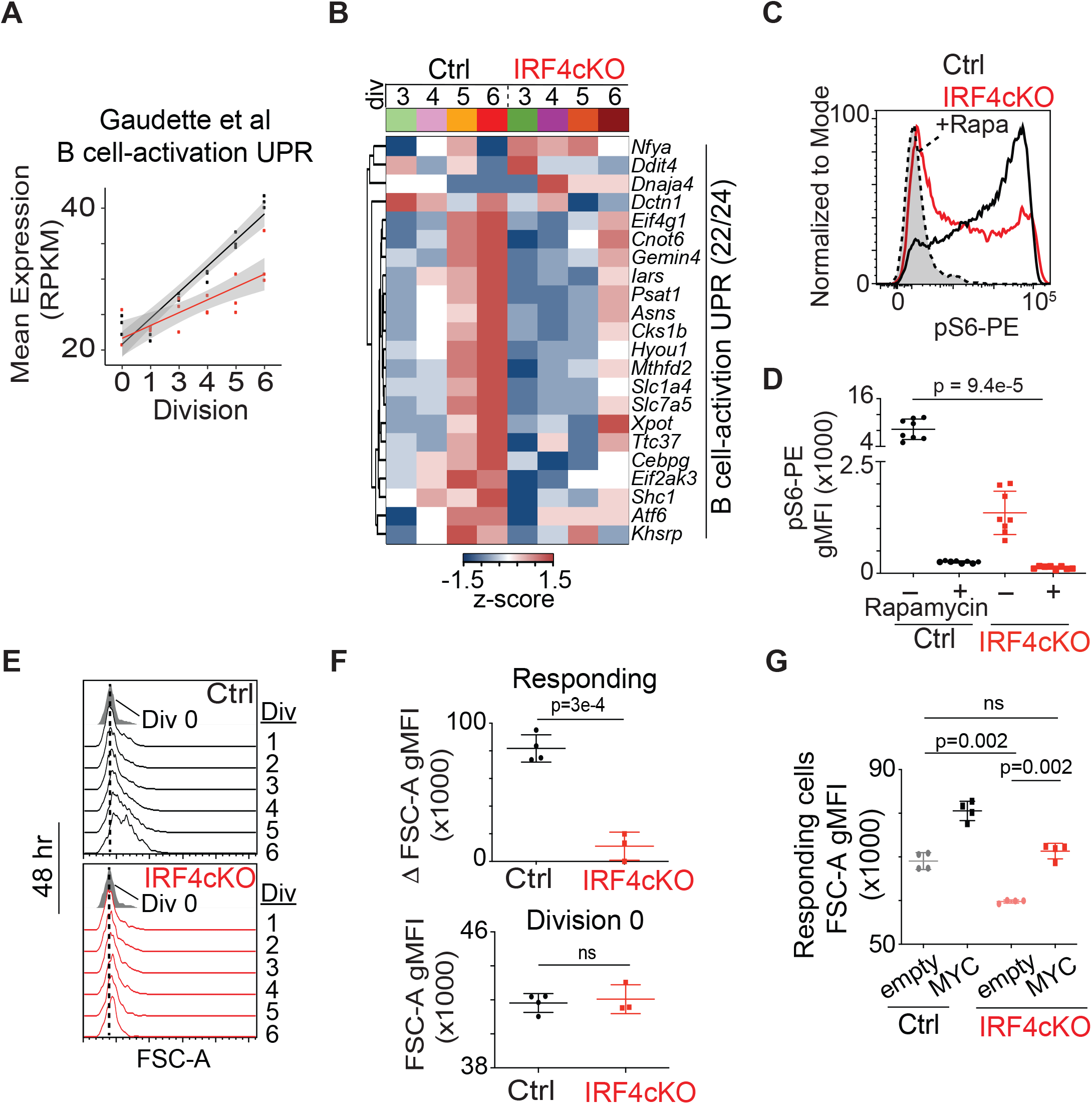
IRF4-deficient B cells exhibit reduced mTORC1 activity and fail to initiate the B cell-activation UPR. **(A)** RNA-seq (described in Fig 4) average RPKM of all detected genes (22/24) in the B cell-activation UPR gene set (31). **(B)** Heatmap of *z* score-normalized gene expression data for all detected genes from **A** for the indicated divisions. **(C)** Representative flow cytometry histograms displaying intracellular phosphorylated S6 (pS6) protein staining for Ctrl or IRF4cKO activated B cells cultured ex vivo with LPS, IL2, and IL5 for 48 h. Grey histogram is representative of Ctrl cultures treated with rapamycin to block mTORC1 activity 2 h before harvest. **(D)** Quantification of geometric mean fluorescence intensity (gMFI) for pS6 from **C. (E)** Histograms displaying cell size distribution via forward scatter area (FSC-A) at divisions 0 - 6 48 h post-LPS inoculation of adoptive transfer host mice, as described in Fig 1. Grey histogram represents cell size at division 0, with the dashed line drawn from the summit to better visualize changes in cell size across the divisions. Cell divisions are indicated to the right of each trace. **(F)** Quantification of data from **E** indicating cell size at division 0 (bottom) and the average change in cell size among responding cells (top). **(G)** Quantification of cell size via forward scatter area (FSC-A) for all responding cells in Ctrl and IRF4cKO transduced with empty-RFP retrovirus or MYC-RFP expressing retrovirus. All data are representative of at least two independent experiments using at least 3 mice per genotype. Data in **D, F**, and **G** represent mean ± SD. Statistical significance in **D** and **F** was determined by a two-tailed Student’s *t* test. Statistical significance in **G** when comparing IRF4cKO samples was determined by a paired two-tailed Student’s *t* test, while significance between Ctrl and IRF4cKO samples was calculated by a two-tailed Student’s *t* test.

## Discussion

This study establishes the timing and extent of IRF4-dependent reprogramming instructed in the initial stages of B cell differentiation in vivo and ascribe a role for IRF4 in controlling cell growth and proliferation. Using multiple antigen model systems, IRF4-deficient B cells divided initially, but stalled during the proliferative response. Characterization of the proliferative defect revealed fewer actively dividing cells and abnormal cell cycle distribution. B cells lacking IRF4 maintained an inflammatory gene signature but failed to induce critical actB and ASC gene expression programs, including metabolic pathways (glycolysis and OXPHOS), MYC target genes, and mTORC1 signaling. Reduced *Myc* expression and mTORC1 activity contributed to the cell division and growth defect following stimulation. Additionally, IRF4-deficient B cells failed to induce the B cell-activation UPR, which relies on mTORC1 (31). Thus, we define the cell division-coupled IRF4-dependent reprogramming events that occur in the initial stages of B cell activation and identify an IRF4-MYC-mTORC1 relationship that impacts cell growth and proliferation.

The role of MYC as a division-independent timer to regulate lymphocyte proliferation has been described (4, 12). In this model, the combination and strength of stimuli determine the amount of MYC initially generated. This serves as a timer to regulate the overall number of cell divisions, or a cell’s division destiny, with division cessation occurring when MYC levels fall below a critical level (12). Analyzing the cell division kinetics of IRF4-deficient B cells responding to LPS revealed they can initiate cell division appropriately but stall in the middle of the proliferative response. Applying the MYC dilution model, IRF4-deficient B cells fall below the MYC threshold sooner, which caused the observed stalling. Indeed, IRF4-deficient B cells displayed reduced MYC levels 24 h after stimulation. Interestingly, MYC expression is not dependent on cell division (12), but we found progressive dysregulation of MYC target genes in IRF4-deficient B cells, implying that other factors reinforce MYC programming throughout the cell divisions. Importantly, both *Irf4* expression and *Myc* induction levels are dependent on the strength of signaling *(12)*, irrespective of whether the stimulus is from BCR (18) or TLR (78). Furthermore, IRF4 binding to the *Myc* promoter has been reported (69, 79). While no differences in chromatin accessibility were observed in IRF4-deficient B cells at known regulatory elements of *Myc* (80), this is likely due to the timing in which the samples were collected or compensatory effects of IRF8 (81, 82), which often binds to the same sites. Collectively, these data support the concept that IRF4 serves as a rheostat in B cells to regulate the overall proliferative response by fine-tuning initial *Myc* expression levels. Indeed, a similar role for IRF4 has been noted in CD8 T cells in which IRF4 serves as a molecular rheostat of TCR affinity. Similar to our observations, IRF4-deficient CD8 T cells can initiate proliferation but fail to maintain clonal expansion (83), suggesting IRF4 may play a similar role in controlling the proliferative response in T cells.

Differentiating actB undergo an IRF4-dependent bifurcation event that commits a portion of actB to an ASC fate (17). Additionally, differentiating actB utilize mTORC1 to anticipate antibody synthesis by upregulating UPR-affiliated genes (31). We demonstrate that IRF4-deficient B cells display reduced mTORC1 activity and fail to initiate the B cell-activation UPR. Thus, actB anticipation of antibody synthesis and secretion is a component of ASC fate commitment and programmed during the initial stages of B cell activation. Our gene expression data indicate that this process occurs as early as division 3 during B cell differentiation, with reduced expression of UPR-affiliated genes in IRF4-deficient B cells. Interestingly, the interplay between mTORC1 and IRF4 has been noted, with mTOR inhibition negatively impacting IRF4 expression (84-86). Here, IRF4 also impacts mTORC1 activity, suggesting the existence of a positive IRF4-mTORC1 feedback loop that impacts actB reprogramming. MYC is central to this regulatory network, as MYC overexpression in IRF4-deficient B cells restores cell growth. mTOR may impact IRF4 transcription by effecting downstream transcription factors or by directly impacting IRF4 protein translation or stability (87).

Occupancy of IRF4 at composite motifs is dependent on its concentration and availability of binding partners (15). IRF4 levels are increased as the cells divide and ultimately sustained at high levels in ASC (17, 18). In contrast, IRF8 levels are decreased as B cells differentiate, allowing for IRF4 to more readily partner with transcription factors and establish the IRF4-dependent gene expression program (82). In IRF4cKO cells, differentiating cells showed changes in accessibility surrounding composite motifs. Previously, ATAC-seq data in wild-type differentiating B cells suggested that EICE motifs were most accessible in early dividing actB and that AICE sites became increasingly accessible as IRF4 levels increased during the division-coupled differentiation process (27). In the absence of IRF4, this program is altered. In regions that decreased accessibility, AICE motifs were the most affected motifs in early divisions (divisions 3 and 4), while EICE motifs were most highly ranked at later divisions (divisions 5 and 6). Although both motifs are affected at all divisions, this analysis pinpoints specific divisions and differentiation stages where IRF4 cooperates with AP-1 or ETS factors to establish differentiation programs, suggesting a hierarchy of IRF4 activity. Consistent with these data, single-cell analysis of LPS responding B cells showed that IRF4 was required for BATF (an AP-1 family member) targets as early as division 3, suggesting that IRF4 may be BATF’s partner in AICEs at the early stages of B cell differentiation to ASC (17). Furthermore, IRF4 binding at AICE motifs largely occurs at newly established accessible regions (14). Taken together, these data indicate that these reprogramming steps occur at divisions 3 and 4.

The cell division requirement needed for ASC formation in vivo following LPS (17, 25, 27) and NP-ficoll (17) stimulation has been described. We observed similar cell division requirements for adoptive transfers using Ctrl B cells and add that ASC formation occurs after cells reach or exceed division 8 following stimulation with the T-dependent antigen influenza X31. As this analysis was performed at day 6 following infection, it is unlikely that the generation of ASC at this time point involve a full germinal center reaction. However, antigen-specific ASC can be observed at this time point (88). These data suggest that the timing of division-coupled reprogramming events needed for ASC differentiation are similar for T-independent antigens and the early differentiation process that occurs with T-dependent antigens. Studying the cell division requirement of T-dependent ASC formation at later time points is complicated by the dynamics and selection pressures of the germinal center reaction and increased cell divisions (89, 90).

Together, these data indicate IRF4 coordinates cell growth and the proliferative response during B cell differentiation. We demonstrate that part of the mechanism involves regulation of *Myc* and mTORC1 activity. Indeed, the relationship between MYC and mTORC1 has been noted, with mTORC1 controlling MYC translation (91) and MYC-driven tumorigenesis dependent on mTORC1 (72, 75). Both factors converge to control protein production and cell growth. MYC controls the expression of translation initiation factors needed for increased protein synthesis (74) and mTOR controls their activity (92). Here, IRF4cKO cells displayed reduced mTORC1 activity and were unable to increase in cell size as they divided. However, the deficiency in cell growth was overcome by overexpression of *Myc*, suggesting that this aspect of MYC/mTOR relationship is dependent on *Myc* expression. RNA-seq analyses showed IRF4-deficient B cells failed to induce MYC target genes and mTORC1 signaling by division 3, and these gene sets became progressively dysregulated as the cells divided. Thus, reprogramming events needed for continued cell growth and proliferation occur during the initial cell divisions during B cell differentiation and are coordinated by IRF4, MYC, and mTORC1.

## Supporting information

Supplemental Figures Patterson et al.

## Acknowledgements

We thank the Boss and Scharer laboratories for their scientific contributions and critical reading of the manuscript, Royce Butler for mouse colony maintenance and husbandry, Tian Mi for bioinformatic assistance, Sakeenah L. Hicks for preparation of sequencing libraries, Dr. Chaoran Li for retroviral reagents (gifted Plat-E cells and pCL-Eco) and protocols, Dr. Troy D. Randall for influenza A/HK-X31 viral stocks, Drs. Susanne Heinzel and Philip Hodgkin for discussions regarding intracellular staining of MYC, the Emory Flow Cytometry Core for FACS isolation of cells, and the Emory Integrated Genetics and Computational Core for Bioanalyzer and sequencing library QC.

## Abbreviations

actB: activated B cell
ASC: antibody-secreting plasma cell
ATAC-seq: assay for transposase accessible chromatin-sequencing
CI: confidence interval
Ctrl: CD45.2+Cd19+/+Irf4fl/fl
CTV: CellTrace Violet
DAR: differentially accessible region
DEG: differentially expressed genes
FDR: false discovery rate
FSC-A: forward scatter area
gMFI: geometric mean fluorescence intensity
GSEA: gene set enrichment analysis
IRF4cKO: CD45.2+Cd19Cre/+Irf4fl/fl
MDN: mean division number
mTOR: mammalian target of rapamycin
nB: naïve B cell
NES: normalized enrichment score
OXPHOS: oxidative phosphorylation
pS6: phosporylated S6
RPKM: reads per kilobase million
scRNA-seq: single cell RNA-sequencing
t-SNE: t-stochastic neighbor embedded
UPR: unfolded protein response

## References

1. Hasbold, J., L. M. Corcoran, D. M. Tarlinton, S. G. Tangye, and P. D. Hodgkin. 2004. Evidence from the generation of immunoglobulin G-secreting cells that stochastic mechanisms regulate lymphocyte differentiation. Nat Immunol 5: 55–63.

2. Hodgkin, P. D., J. H. Lee, and A. B. Lyons. 1996. B cell differentiation and isotype switching is related to division cycle number. J Exp Med 184: 277–281.

3. Jelinek, D. F., and P. E. Lipsky. 1983. The role of B cell proliferation in the generation of immunoglobulin-secreting cells in man. J Immunol 130: 2597–2604.

4. Heinzel, S., J. M. Marchingo, M. B. Horton, and P. D. Hodgkin. 2018. The regulation of lymphocyte activation and proliferation. Curr Opin Immunol 51: 32–38.

5. Taylor, J. J., K. A. Pape, H. R. Steach, and M. K. Jenkins. 2015. Humoral immunity. Apoptosis and antigen affinity limit effector cell differentiation of a single naive B cell. Science 347: 784–787.

6. Duffy, K. R., C. J. Wellard, J. F. Markham, J. H. Zhou, R. Holmberg, E. D. Hawkins, J. Hasbold, M. R. Dowling, and P. D. Hodgkin. 2012. Activation-induced B cell fates are selected by intracellular stochastic competition. Science 335: 338–341.

7. Zhou, J. H. S., J. F. Markham, K. R. Duffy, and P. D. Hodgkin. 2018. Stochastically Timed Competition Between Division and Differentiation Fates Regulates the Transition From B Lymphoblast to Plasma Cell. Frontiers in immunology 9: 2053.

8. Mitchell, S. 2020. What Will B Will B: Identifying Molecular Determinants of Diverse B-Cell Fate Decisions Through Systems Biology. Front Cell Dev Biol 8: 616592.

9. Fernandez, D., M. Ortiz, L. Rodriguez, A. Garcia, D. Martinez, and I. Moreno de Alboran. 2013. The proto-oncogene c-myc regulates antibody secretion and Ig class switch recombination. J Immunol 190: 6135–6144.

10. Finkin, S., H. Hartweger, T. Y. Oliveira, E. E. Kara, and M. C. Nussenzweig. 2019. Protein Amounts of the MYC Transcription Factor Determine Germinal Center B Cell Division Capacity. Immunity.

11. Perez-Olivares, M., A. Trento, S. Rodriguez-Acebes, D. Gonzalez-Acosta, D. Fernandez-Antoran, S. Roman-Garcia, D. Martinez, T. Lopez-Briones, C. Torroja, Y. R. Carrasco, J. Mendez, and I. Moreno de Alboran. 2018. Functional interplay between c-Myc and Max in B lymphocyte differentiation. EMBO Rep 19.

12. Heinzel, S., T. Binh Giang, A. Kan, J. M. Marchingo, B. K. Lye, L. M. Corcoran, and P. D. Hodgkin. 2017. A Myc-dependent division timer complements a cell-death timer to regulate T cell and B cell responses. Nat Immunol 18: 96–103.

13. Hawkins, E. D., M. L. Turner, C. J. Wellard, J. H. Zhou, M. R. Dowling, and P. D. Hodgkin. 2013. Quantal and graded stimulation of B lymphocytes as alternative strategies for regulating adaptive immune responses. Nat Commun 4: 2406.

14. Ochiai, K., M. Maienschein-Cline, G. Simonetti, J. Chen, R. Rosenthal, R. Brink, A. S. Chong, U. Klein, A. R. Dinner, H. Singh, and R. Sciammas. 2013. Transcriptional regulation of germinal center B and plasma cell fates by dynamical control of IRF4. Immunity 38: 918–929.

15. Sciammas, R., A. L. Shaffer, J. H. Schatz, H. Zhao, L. M. Staudt, and H. Singh. 2006. Graded expression of interferon regulatory factor-4 coordinates isotype switching with plasma cell differentiation. Immunity 25: 225–236.

16. Klein, U., S. Casola, G. Cattoretti, Q. Shen, M. Lia, T. Mo, T. Ludwig, K. Rajewsky, and R. Dalla-Favera. 2006. Transcription factor IRF4 controls plasma cell differentiation and class-switch recombination. Nat Immunol 7: 773–782.

17. Scharer, C. D., D. G. Patterson, T. Mi, M. J. Price, S. L. Hicks, and J. M. Boss. 2020. Antibody-secreting cell destiny emerges during the initial stages of B-cell activation. Nat Commun 11: 3989.

18. Sciammas, R., Y. Li, A. Warmflash, Y. Song, A. R. Dinner, and H. Singh. 2011. An incoherent regulatory network architecture that orchestrates B cell diversification in response to antigen signaling. Mol Syst Biol 7: 495.

19. Mittrucker, H. W., T. Matsuyama, A. Grossman, T. M. Kundig, J. Potter, A. Shahinian, Wakeham, B. Patterson, P. S. Ohashi, and T. W. Mak. 1997. Requirement for the transcription factor LSIRF/IRF4 for mature B and T lymphocyte function. Science 275: 540–543.

20. Di Pietro, A., and K. L. Good-Jacobson. 2018. Disrupting the Code: Epigenetic Dysregulation of Lymphocyte Function during Infectious Disease and Lymphoma Development. J Immunol 201: 1109–1118.

21. Haines, R. R., B. G. Barwick, C. D. Scharer, P. Majumder, T. D. Randall, and J. M. Boss. 2018. The Histone Demethylase LSD1 Regulates B Cell Proliferation and Plasmablast Differentiation. J Immunol 201: 2799–2811.

22. Barwick, B. G., C. D. Scharer, R. J. Martinez, M. J. Price, A. N. Wein, R. R. Haines, A. P. R. Bally, J. E. Kohlmeier, and J. M. Boss. 2018. B cell activation and plasma cell differentiation are inhibited by de novo DNA methylation. Nat Commun 9: 1900.

23. Guo, M., M. J. Price, D. G. Patterson, B. G. Barwick, R. R. Haines, A. K. Kania, J. E. Bradley, T. D. Randall, J. M. Boss, and C. D. Scharer. 2018. EZH2 Represses the B Cell Transcriptional Program and Regulates Antibody-Secreting Cell Metabolism and Antibody Production. J Immunol 200: 1039–1052.

24. Price, M. J., D. G. Patterson, C. D. Scharer, and J. M. Boss. 2018. Progressive Upregulation of Oxidative Metabolism Facilitates Plasmablast Differentiation to a T-Independent Antigen. Cell Rep 23: 3152–3159.

25. Barwick, B. G., C. D. Scharer, A. P. R. Bally, and J. M. Boss. 2016. Plasma cell differentiation is coupled to division-dependent DNA hypomethylation and gene regulation. Nat Immunol 17: 1216–1225.

26. Wiggins, K. J., and C. D. Scharer. 2021. Roadmap to a plasma cell: Epigenetic and transcriptional cues that guide B cell differentiation. Immunol Rev 300: 54–64.

27. Scharer, C. D., B. G. Barwick, M. Guo, A. P. R. Bally, and J. M. Boss. 2018. Plasma cell differentiation is controlled by multiple cell division-coupled epigenetic programs. Nat Commun 9: 1698.

28. Nutt, S. L., P. D. Hodgkin, D. M. Tarlinton, and L. M. Corcoran. 2015. The generation of antibody-secreting plasma cells. Nat Rev Immunol 15: 160–171.

29. Walter, P., and D. Ron. 2011. The unfolded protein response: from stress pathway to homeostatic regulation. Science 334: 1081–1086.

30. Bettigole, S. E., and L. H. Glimcher. 2015. Endoplasmic reticulum stress in immunity. Annu Rev Immunol 33: 107–138.

31. Gaudette, B. T., D. D. Jones, A. Bortnick, Y. Argon, and D. Allman. 2020. mTORC1 coordinates an immediate unfolded protein response-related transcriptome in activated B cells preceding antibody secretion. Nat Commun 11: 723.

32. Iwakoshi, N. N., A. H. Lee, and L. H. Glimcher. 2003. The X-box binding protein-1 transcription factor is required for plasma cell differentiation and the unfolded protein response. Immunol Rev 194: 29–38.

33. Shaffer, A. L., M. Shapiro-Shelef, N. N. Iwakoshi, A. H. Lee, S. B. Qian, H. Zhao, X. Yu, L. Yang, B. K. Tan, A. Rosenwald, E. M. Hurt, E. Petroulakis, N. Sonenberg, J. W. Yewdell, K. Calame, L. H. Glimcher, and L. M. Staudt. 2004. XBP1, downstream of Blimp-1, expands the secretory apparatus and other organelles, and increases protein synthesis in plasma cell differentiation. Immunity 21: 81–93.

34. Buenrostro, J. D., B. Wu, H. Y. Chang, and W. J. Greenleaf. 2015. ATAC-seq: A Method for Assaying Chromatin Accessibility Genome-Wide. Curr Protoc Mol Biol 109: 21 29 21–29.

35. Scharer, C. D., E. L. Blalock, B. G. Barwick, R. R. Haines, C. Wei, I. Sanz, and J. M. Boss. 2016. ATAC-seq on biobanked specimens defines a unique chromatin accessibility structure in naive SLE B cells. Sci Rep 6: 27030.

36. Rickert, R. C., J. Roes, and K. Rajewsky. 1997. B lymphocyte-specific, Cre-mediated mutagenesis in mice. Nucleic Acids Res 25: 1317–1318.

37. Kitamura, D., J. Roes, R. Kuhn, and K. Rajewsky. 1991. A B cell-deficient mouse by targeted disruption of the membrane exon of the immunoglobulin mu chain gene. Nature 350: 423–426.

38. Yoon, H. S., C. D. Scharer, P. Majumder, C. W. Davis, R. Butler, W. Zinzow-Kramer, I. Skountzou, D. G. Koutsonanos, R. Ahmed, and J. M. Boss. 2012. ZBTB32 is an early repressor of the CIITA and MHC class II gene expression during B cell differentiation to plasma cells. J Immunol 189: 2393–2403.

39. Magnuson, A. M., E. Kiner, A. Ergun, J. S. Park, N. Asinovski, A. Ortiz-Lopez, A. Kilcoyne, E. Paoluzzi-Tomada, R. Weissleder, D. Mathis, and C. Benoist. 2018. Identification and validation of a tumor-infiltrating Treg transcriptional signature conserved across species and tumor types. Proc Natl Acad Sci U S A 115: E10672–E10681.

40. Naviaux, R. K., E. Costanzi, M. Haas, and I. M. Verma. 1996. The pCL vector system: rapid production of helper-free, high-titer, recombinant retroviruses. J Virol 70: 5701–5705.

41. Kawauchi, D., G. Robinson, T. Uziel, P. Gibson, J. Rehg, C. Gao, D. Finkelstein, C. Qu, S. Pounds, D. W. Ellison, R. J. Gilbertson, and M. F. Roussel. 2012. A mouse model of the most aggressive subgroup of human medulloblastoma. Cancer Cell 21: 168–180.

42. Dobin, A., C. A. Davis, F. Schlesinger, J. Drenkow, C. Zaleski, S. Jha, P. Batut, M. Chaisson, and T. R. Gingeras. 2013. STAR: ultrafast universal RNA-seq aligner. Bioinformatics 29: 15–21.

43. Robinson, M. D., D. J. McCarthy, and G. K. Smyth. 2010. edgeR: a Bioconductor package for differential expression analysis of digital gene expression data. Bioinformatics 26: 139–140.

44. Subramanian, A., P. Tamayo, V. K. Mootha, S. Mukherjee, B. L. Ebert, M. A. Gillette, Paulovich, S. L. Pomeroy, T. R. Golub, E. S. Lander, and J. P. Mesirov. 2005. Gene set enrichment analysis: a knowledge-based approach for interpreting genome-wide expression profiles. Proc Natl Acad Sci U S A 102: 15545–15550.

45. Langmead, B., C. Trapnell, M. Pop, and S. L. Salzberg. 2009. Ultrafast and memory-efficient alignment of short DNA sequences to the human genome. Genome Biol 10: R25.

46. Zhang, Y., T. Liu, C. A. Meyer, J. Eeckhoute, D. S. Johnson, B. E. Bernstein, C. Nusbaum, R. M. Myers, M. Brown, W. Li, and X. S. Liu. 2008. Model-based analysis of ChIP-Seq (MACS). Genome Biol 9: R137.

47. Heinz, S., C. Benner, N. Spann, E. Bertolino, Y. C. Lin, P. Laslo, J. X. Cheng, C. Murre, H. Singh, and C. K. Glass. 2010. Simple combinations of lineage-determining transcription factors prime cis-regulatory elements required for macrophage and B cell identities. Mol Cell 38: 576–589.

48. Smith, K. G., T. D. Hewitson, G. J. Nossal, and D. M. Tarlinton. 1996. The phenotype and fate of the antibody-forming cells of the splenic foci. Eur J Immunol 26: 444–448.

49. Kallies, A., J. Hasbold, D. M. Tarlinton, W. Dietrich, L. M. Corcoran, P. D. Hodgkin, and S. L. Nutt. 2004. Plasma cell ontogeny defined by quantitative changes in blimp-1 expression. J Exp Med 200: 967–977.

50. Turner, M. L., E. D. Hawkins, and P. D. Hodgkin. 2008. Quantitative regulation of B cell division destiny by signal strength. J Immunol 181: 374–382.

51. Vignon, C., C. Debeissat, M. T. Georget, D. Bouscary, E. Gyan, P. Rosset, and O. Herault. 2013. Flow cytometric quantification of all phases of the cell cycle and apoptosis in a two-color fluorescence plot. PLoS One 8: e68425.

52. Lam, W. Y., and D. Bhattacharya. 2018. Metabolic Links between Plasma Cell Survival, Secretion, and Stress. Trends Immunol 39: 19–27.

53. Jin, L., J. Chun, C. Pan, G. N. Alesi, D. Li, K. R. Magliocca, Y. Kang, Z. G. Chen, D. M. Shin, F. R. Khuri, J. Fan, and S. Kang. 2017. Phosphorylation-mediated activation of LDHA promotes cancer cell invasion and tumour metastasis. Oncogene 36: 3797–3806.

54. Chang, Y. C., Y. C. Yang, C. P. Tien, C. J. Yang, and M. Hsiao. 2018. Roles of Aldolase Family Genes in Human Cancers and Diseases. Trends Endocrinol Metab 29: 549–559.

55. Xiong, Y., J. Lu, Q. Fang, Y. Lu, C. Xie, H. Wu, and Z. Yin. 2019. UBE2C functions as a potential oncogene by enhancing cell proliferation, migration, invasion, and drug resistance in hepatocellular carcinoma cells. Biosci Rep 39.

56. Huang, L., H. Y. Wang, J. D. Li, J. H. Wang, Y. Zhou, R. Z. Luo, J. P. Yun, Y. Zhang, W. H. Jia, and M. Zheng. 2013. KPNA2 promotes cell proliferation and tumorigenicity in epithelial ovarian carcinoma through upregulation of c-Myc and downregulation of FOXO3a. Cell Death Dis 4: e745.

57. Gao, Z., X. Man, Z. Li, J. Bi, X. Liu, Z. Li, J. Li, Z. Zhang, and C. Kong. 2020. PLK1 promotes proliferation and suppresses apoptosis of renal cell carcinoma cells by phosphorylating MCM3. Cancer Gene Ther 27: 412–423.

58. Zhu, J., K. Cui, Y. Cui, C. Ma, and Z. Zhang. 2020. PLK1 Knockdown Inhibits Cell Proliferation and Cell Apoptosis, and PLK1 Is Negatively Regulated by miR-4779 in Osteosarcoma Cells. DNA Cell Biol 39: 747–755.

59. Heikkila, R., G. Schwab, E. Wickstrom, S. L. Loke, D. H. Pluznik, R. Watt, and L. M. Neckers. 1987. A c-myc antisense oligodeoxynucleotide inhibits entry into S phase but not progress from G0 to G1. Nature 328: 445–449.

60. Wickstrom, E. L., T. A. Bacon, A. Gonzalez, D. L. Freeman, G. H. Lyman, and E. Wickstrom. 1988. Human promyelocytic leukemia HL-60 cell proliferation and c-myc protein expression are inhibited by an antisense pentadecadeoxynucleotide targeted against c-myc mRNA. Proc Natl Acad Sci U S A 85: 1028–1032.

61. Bretones, G., M. D. Delgado, and J. Leon. 2015. Myc and cell cycle control. Biochim Biophys Acta 1849: 506–516.

62. Buenrostro, J. D., P. G. Giresi, L. C. Zaba, H. Y. Chang, and W. J. Greenleaf. 2013. Transposition of native chromatin for fast and sensitive epigenomic profiling of open chromatin, DNA-binding proteins and nucleosome position. Nat Methods 10: 1213–1218.

63. Krishnamoorthy, V., S. Kannanganat, M. Maienschein-Cline, S. L. Cook, J. Chen, N. Bahroos, E. Sievert, E. Corse, A. Chong, and R. Sciammas. 2017. The IRF4 Gene Regulatory Module Functions as a Read-Write Integrator to Dynamically Coordinate T Helper Cell Fate. Immunity 47: 481–497 e487.

64. Glasmacher, E., S. Agrawal, A. B. Chang, T. L. Murphy, W. Zeng, B. Vander Lugt, A. Khan, M. Ciofani, C. J. Spooner, S. Rutz, J. Hackney, R. Nurieva, C. R. Escalante, W. Ouyang, D. R. Littman, K. M. Murphy, and H. Singh. 2012. A genomic regulatory element that directs assembly and function of immune-specific AP-1-IRF complexes. Science 338: 975–980.

65. Brass, A. L., A. Q. Zhu, and H. Singh. 1999. Assembly requirements of PU.1-Pip (IRF-4) activator complexes: inhibiting function in vivo using fused dimers. EMBO J 18: 977–991.

66. Eisenbeis, C. F., H. Singh, and U. Storb. 1995. Pip, a novel IRF family member, is a lymphoid-specific, PU.1-dependent transcriptional activator. Genes Dev 9: 1377–1387.

67. Ochiai, K., Y. Katoh, T. Ikura, Y. Hoshikawa, T. Noda, H. Karasuyama, S. Tashiro, A. Muto, and K. Igarashi. 2006. Plasmacytic transcription factor Blimp-1 is repressed by Bach2 in B cells. J Biol Chem 281: 38226–38234.

68. Stone, S. L., J. N. Peel, C. D. Scharer, C. A. Risley, D. A. Chisolm, M. D. Schultz, B. Yu, Ballesteros-Tato, W. Wojciechowski, B. Mousseau, R. S. Misra, A. Hanidu, H. Jiang, Z. Qi, J. M. Boss, T. D. Randall, S. R. Brodeur, A. W. Goldrath, A. S. Weinmann, A. F. Rosenberg, and F. E. Lund. 2019. T-bet Transcription Factor Promotes Antibody-Secreting Cell Differentiation by Limiting the Inflammatory Effects of IFN-gamma on B Cells. Immunity 50: 1172–1187 e1177.

69. Shaffer, A. L., N. C. Emre, L. Lamy, V. N. Ngo, G. Wright, W. Xiao, J. Powell, S. Dave, X. Yu, H. Zhao, Y. Zeng, B. Chen, J. Epstein, and L. M. Staudt. 2008. IRF4 addiction in multiple myeloma. Nature 454: 226–231.

70. Hann, S. R. 2006. Role of post-translational modifications in regulating c-Myc proteolysis, transcriptional activity and biological function. Semin Cancer Biol 16: 288–302.

71. Saxton, R. A., and D. M. Sabatini. 2017. mTOR Signaling in Growth, Metabolism, and Disease. Cell 169: 361–371.

72. Liu, P., M. Ge, J. Hu, X. Li, L. Che, K. Sun, L. Cheng, Y. Huang, M. G. Pilo, A. Cigliano, G. M. Pes, R. M. Pascale, S. Brozzetti, G. Vidili, A. Porcu, A. Cossu, G. Palmieri, M. C. Sini, S. Ribback, F. Dombrowski, J. Tao, D. F. Calvisi, L. Chen, and X. Chen. 2017. A functional mammalian target of rapamycin complex 1 signaling is indispensable for c-Myc-driven hepatocarcinogenesis. Hepatology 66: 167–181.

73. Gao, S., M. Chen, W. Wei, X. Zhang, M. Zhang, Y. Yao, Y. Lv, T. Ling, L. Wang, and X. Zou. 2018. Crosstalk of mTOR/PKM2 and STAT3/c-Myc signaling pathways regulate the energy metabolism and acidic microenvironment of gastric cancer. J Cell Biochem.

74. Lin, C. J., R. Cencic, J. R. Mills, F. Robert, and J. Pelletier. 2008. c-Myc and eIF4F are components of a feedforward loop that links transcription and translation. Cancer Res 68: 5326–5334.

75. Pourdehnad, M., M. L. Truitt, I. N. Siddiqi, G. S. Ducker, K. M. Shokat, and D. Ruggero. 2013. Myc and mTOR converge on a common node in protein synthesis control that confers synthetic lethality in Myc-driven cancers. Proc Natl Acad Sci U S A 110: 11988–11993.

76. Schmidt, E. V., M. J. Ravitz, L. Chen, and M. Lynch. 2009. Growth controls connect: interactions between c-myc and the tuberous sclerosis complex-mTOR pathway. Cell Cycle 8: 1344–1351.

77. Sriburi, R., S. Jackowski, K. Mori, and J. W. Brewer. 2004. XBP1: a link between the unfolded protein response, lipid biosynthesis, and biogenesis of the endoplasmic reticulum. J Cell Biol 167: 35–41.

78. Negishi, H., Y. Ohba, H. Yanai, A. Takaoka, K. Honma, K. Yui, T. Matsuyama, T. Taniguchi, and K. Honda. 2005. Negative regulation of Toll-like-receptor signaling by IRF-4. Proc Natl Acad Sci U S A 102: 15989–15994.

79. Boddicker, R. L., N. S. Kip, X. Xing, Y. Zeng, Z. Z. Yang, J. H. Lee, L. L. Almada, S. F. Elsawa, R. A. Knudson, M. E. Law, R. P. Ketterling, J. M. Cunningham, Y. Wu, M. J. Maurer, M. M. O’Byrne, J. R. Cerhan, S. L. Slager, B. K. Link, J. C. Porcher, D. M. Grote, D. F. Jelinek, A. Dogan, S. M. Ansell, M. E. Fernandez-Zapico, and A. L. Feldman. 2015. The oncogenic transcription factor IRF4 is regulated by a novel CD30/NF-kappaB positive feedback loop in peripheral T-cell lymphoma. Blood 125: 3118–3127.

80. Chaudhri, V. K., K. Dienger-Stambaugh, Z. Wu, M. Shrestha, and H. Singh. 2020. Charting the cis-regulome of activated B cells by coupling structural and functional genomics. Nat Immunol 21: 210–220.

81. Carotta, S., S. N. Willis, J. Hasbold, M. Inouye, S. H. Pang, D. Emslie, A. Light, M. Chopin, W. Shi, H. Wang, H. C. Morse, 3rd, D. M. Tarlinton, L. M. Corcoran, P. D. Hodgkin, and S. L. Nutt. 2014. The transcription factors IRF8 and PU.1 negatively regulate plasma cell differentiation. J Exp Med 211: 2169–2181.

82. Xu, H., V. K. Chaudhri, Z. Wu, K. Biliouris, K. Dienger-Stambaugh, Y. Rochman, and H. Singh. 2015. Regulation of bifurcating B cell trajectories by mutual antagonism between transcription factors IRF4 and IRF8. Nat Immunol 16: 1274–1281.

83. Man, K., M. Miasari, W. Shi, A. Xin, D. C. Henstridge, S. Preston, M. Pellegrini, G. T. Belz, G. K. Smyth, M. A. Febbraio, S. L. Nutt, and A. Kallies. 2013. The transcription factor IRF4 is essential for TCR affinity-mediated metabolic programming and clonal expansion of T cells. Nat Immunol 14: 1155–1165.

84. Chapman, N. M., H. Zeng, T. M. Nguyen, Y. Wang, P. Vogel, Y. Dhungana, X. Liu, G. Neale, J. W. Locasale, and H. Chi. 2018. mTOR coordinates transcriptional programs and mitochondrial metabolism of activated Treg subsets to protect tissue homeostasis. Nat Commun 9: 2095.

85. Raybuck, A. L., S. H. Cho, J. Li, M. C. Rogers, K. Lee, C. L. Williams, M. Shlomchik, J. W. Thomas, J. Chen, J. V. Williams, and M. R. Boothby. 2018. B Cell-Intrinsic mTORC1 Promotes Germinal Center-Defining Transcription Factor Gene Expression, Somatic Hypermutation, and Memory B Cell Generation in Humoral Immunity. J Immunol 200: 2627–2639.

86. Yao, S., B. F. Buzo, D. Pham, L. Jiang, E. J. Taparowsky, M. H. Kaplan, and J. Sun. 2013. Interferon regulatory factor 4 sustains CD8(+) T cell expansion and effector differentiation. Immunity 39: 833–845.

87. Laplante, M., and D. M. Sabatini. 2013. Regulation of mTORC1 and its impact on gene expression at a glance. J Cell Sci 126: 1713–1719.

88. Price, M. J., C. D. Scharer, A. K. Kania, T. D. Randall, and J. M. Boss. 2021. Conserved Epigenetic Programming and Enhanced Heme Metabolism Drive Memory B Cell Reactivation. J Immunol.

89. Victora, G. D., and M. C. Nussenzweig. 2012. Germinal centers. Annu Rev Immunol 30: 429–457.

90. Mesin, L., J. Ersching, and G. D. Victora. 2016. Germinal Center B Cell Dynamics. Immunity 45: 471–482.

91. Wall, M., G. Poortinga, K. M. Hannan, R. B. Pearson, R. D. Hannan, and G. A. McArthur. 2008. Translational control of c-MYC by rapamycin promotes terminal myeloid differentiation. Blood 112: 2305–2317.

92. Hay, N., and N. Sonenberg. 2004. Upstream and downstream of mTOR. Genes Dev 18: 1926–1945.

93. Minnich, M., H. Tagoh, P. Bonelt, E. Axelsson, M. Fischer, B. Cebolla, A. Tarakhovsky, S. L. Nutt, M. Jaritz, and M. Busslinger. 2016. Multifunctional role of the transcription factor Blimp-1 in coordinating plasma cell differentiation. Nat Immunol 17: 331–343.

