## Supplemental Figures Patterson et al. for "An IRF4-MYC-mTORC1 integrated pathway controls cell growth and the proliferative capacity of activated B cells during B cell differentiation *in vivo*"

Supplemental Figure 1

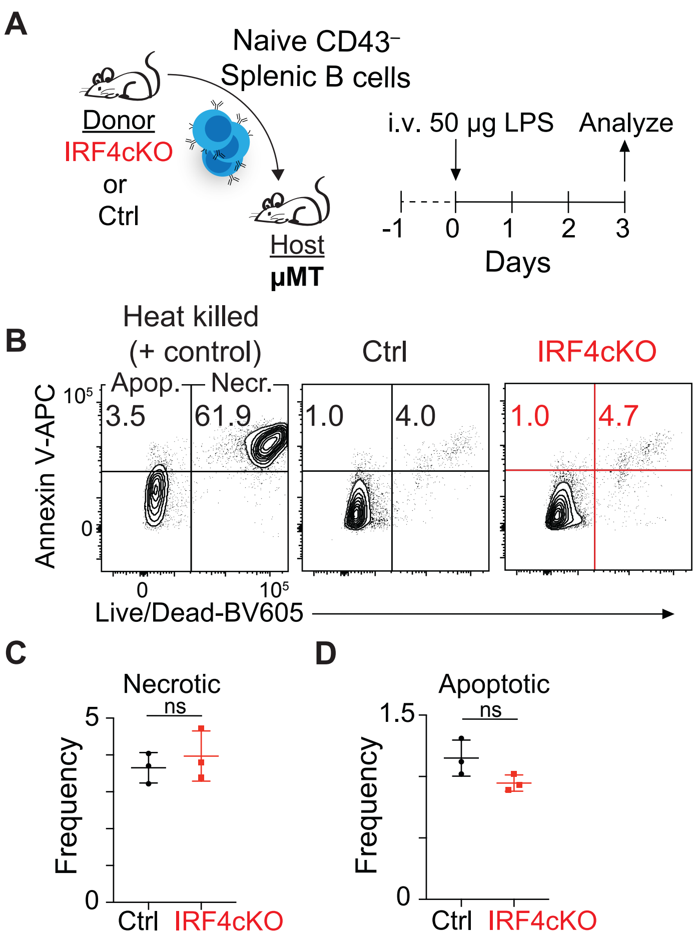


**SUPPLEMENTAL FIGURE 1**. **IRF4-deficient and -sufficient B cells exhibit similar frequencies of apoptosis in vivo.** (**A**) Schematic of experimental design. Ctrl or IRF4cKO cells were transferred into µMT hosts. After 24 h, host mice were challenged with 50 µg LPS and apoptosis was assessed 72 h post-LPS inoculation. (**B**) Representative flow cytometry plots of Annexin V versus Live/Dead viability dye for Ctrl (middle) or IRF4cKO (right). Heat killed control samples were prepared to faithfully gate apoptotic (Apop.) and necrotic (Necr.) cells (left). (**C** and **D**) Frequency of necrotic (C) and apoptotic cells (D) from **B**. Data are representative of two independent experiments using 3 mice per genotype. Statistical significance in **C** and **D** was determined by a two-tailed Student's *t* test.

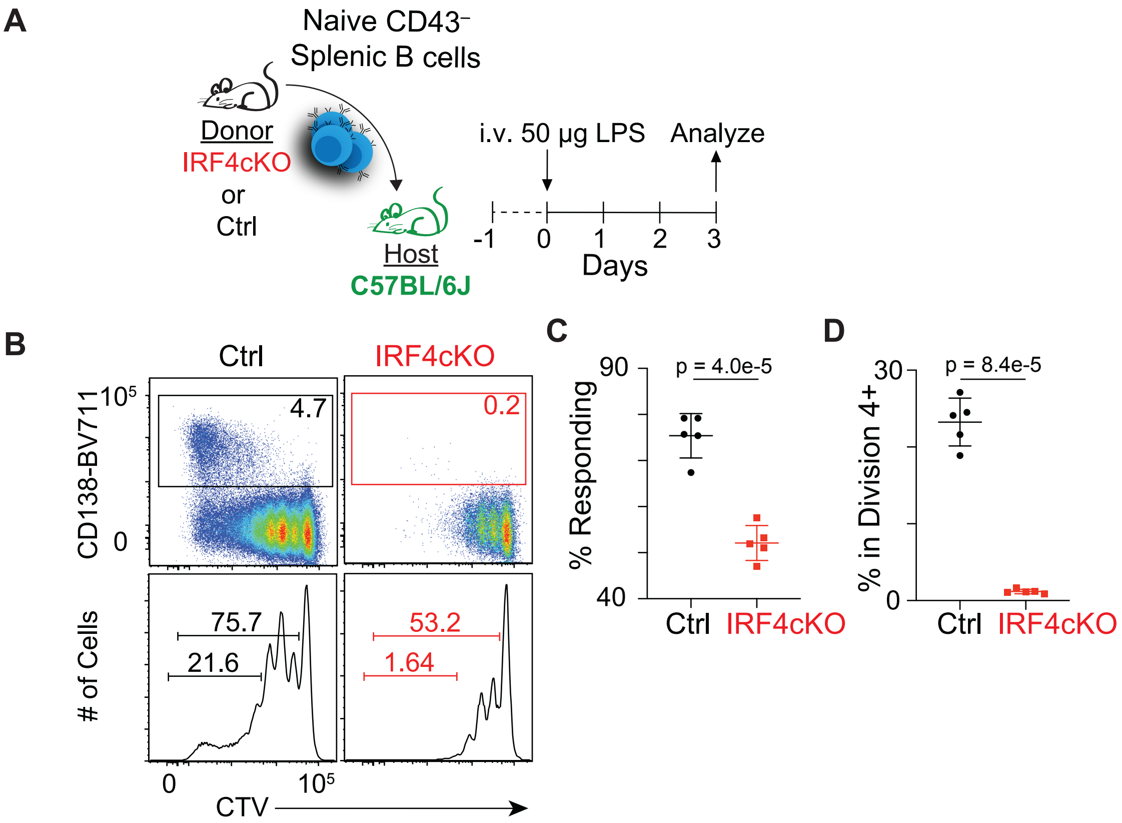


**SUPPLEMENTAL FIGURE 2. IRF4-deficient B cells exhibit a proliferation defect when transferred to C57BL/6J animals.** (**A**) Schematic of experimental design. CTV-labeled Ctrl or IRF4cKO cells were transferred into CD45.1 C57BL/6J hosts. After 24 h, host mice were challenged with 50 µg LPS. Cell division and differentiation were assessed 72 h post-LPS inoculation. (**B**) Representative flow cytometry plots of CD138 versus CTV (top) and CTV histograms (bottom). Frequency of CD138+ ASC are indicated (top), and the frequency of all responding cells, as well as the frequency of cells in divisions 4+ are displayed (bottom). (**C** and **D**) Frequency of all responding cells (C) and cells in division 4+ (D) from **B**. Data are representative of two independent experiments using 3 mice per genotype. Statistical significance in **C** and **D** was determined by a two-tailed Student's *t* test.

Supplemental Figure 3

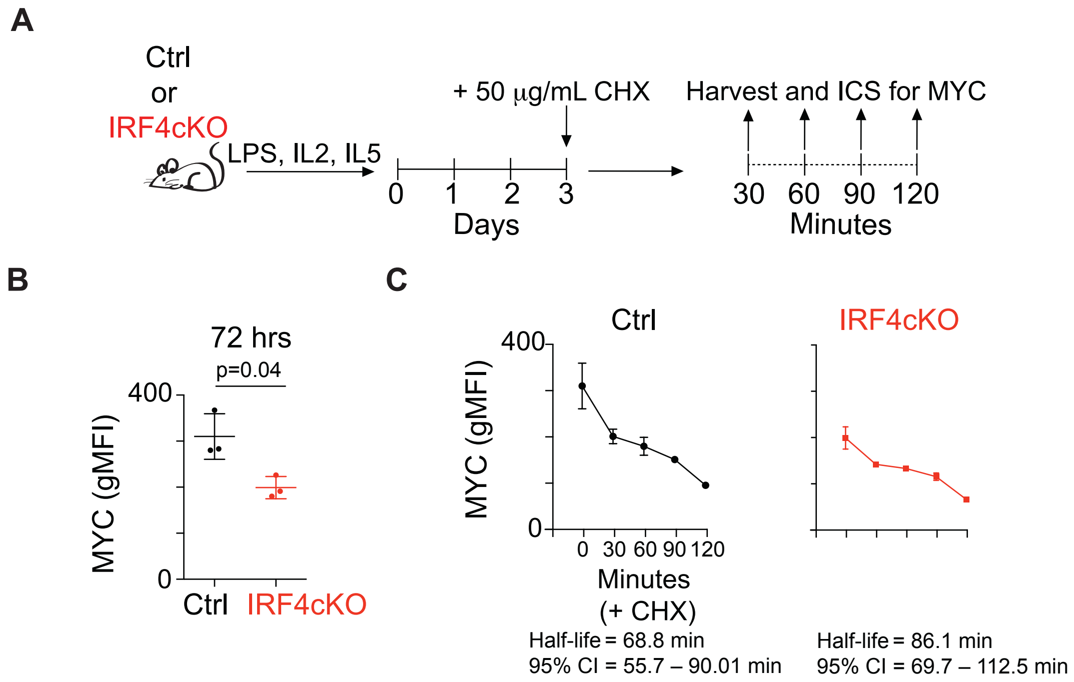


**SUPPLEMENTAL FIGURE 3. MYC protein stability is similar between IRF4-deficient and -sufficient B cells.** (**A**) Schematic of experimental design. Ctrl and IRF4cKO were cultured for 3 days. At 72 h, 50 µg/ml cycloheximide (CHX) was added, and intracellular staining of MYC was performed at the time course shown. (**B**) MYC geometric mean fluorescence intensity (gMFI) at 72 h. (**C**) MYC protein decay curves displayed as gMFI at the indicated time points after CHX treatment. Half-life and 95% confidence intervals (CI) are shown below and were estimated by fitting exponential decay curves. Data are representative of two independent experiments using 3 mice per genotype. Statistical significance in **B** was determined by a two-tailed Student's *t* test.
